# *QCR7* affects the virulence of *Candida albicans* and the uptake of multiple carbon sources present in different host niches

**DOI:** 10.1101/2022.11.02.514977

**Authors:** Lingbing Zeng, Yongcheng Huang, Junjun Tan, Jun Peng, Niya Hu, Qiong Liu, YanLi Cao, Yuping Zhang, Junzhu Chen, Xiaotian Huang

**Affiliations:** The First Affiliated Hospital of Nanchang University, School of Public Health, Jiangxi Medical College, Nanchang University, China; Department of Medical Microbiology, Jiangxi Medical College, Nanchang University, China; School of Public Health, Jiangxi Medical College, Nanchang University/Jiangxi Provincial Key Laboratory of Preventive Medicine, Nanchang University, Nanchang 330006, P.R. China

**Keywords:** *Candida albicans*, virulence, biofilm, hyphae, mitochondrial disorder

## Abstract

**Background:** *Candida albicans* is a commensal yeast that may cause life-threatening infections. Studies have shown that the cytochrome b-c1 complex subunit 7 gene (*QCR7*) of *C. albicans* encodes a protein that forms a component of the mitochondrial electron transport chain complex III, making it an important target for studying the virulence of this yeast. However, to the best of our knowledge, the functions of *QCR7* have not yet been characterized.

**Methods:** A *QCR7* knockout strain was constructed using SN152, and BALb/c mice were used as model animals to determine the role of *QCR7* in the virulence of *C. albicans*. Subsequently, the effects of *QCR7* on mitochondrial functions and use of carbon sources were investigated. Next, its mutant biofilm formation and hyphal growth maintenance were compared with those of the wild type. Furthermore, the transcriptome of the *qcr7Δ/Δ* mutant was compared with that of the WT strain to explore pathogenic mechanisms.

**Results:** Defective *QCR7* attenuated the virulence of *C. albicans* infection in vivo. Furthermore, the mutant influenced the use of multiple alternative carbon sources that exist in several host niches. Moreover, it led to mitochondrial dysfunction. Furthermore, the *QCR7* knockout strain showed defects in biofilm formation or the maintenance of filamentous growth. The overexpression of cell-surface-associated genes (*HWP1, YWP1, XOG1*, and *SAP6*) can restore defective virulence phenotypes and the carbon-source utilization of *qcr7Δ/Δ*.

**Conclusion:** This study provides new insights into the mitochondria-based metabolism of *C. albicans*, accounting for its virulence and the use of variable carbon sources that promote *C. albicans* to colonize host niches.

## 1. Introduction

*Candida albicans* is one of the most important opportunistic pathogens worldwide. Its effects range from superficial to invasive infections and even death. The mortality rate of *C. albicans* during nosocomial infection can be as high as 50% (Gow and Yadav, 2017). The use of various carbon sources is a key factor that influences the virulence of *C. albicans* (Williams and Lorenz, 2020). Although glucose is the preferred carbon source of *C. albicans*, this organism inhabits host environments with limited glucose but rich alternative-carbon sources. Hence, several cellular complexes have evolved in *C. albicans* to use multiple sources of carbon, contributing to its growth and virulence (Brown et al., 2007).

The mitochondrial electron transport chain (METC) is the primary site of carbon metabolism. METC comprises five complexes; most studies only focused on the function of complexes I and V (Huang et al., 2017; Li et al., 2017; Li et al., 2018). However, limited studies have explored complex III. We selected this complex because of its importance in electron transport. Complex III, also known as cytochrome bc1 complex, is a component of the mitochondrial respiratory chain (Tzagoloff, 1995). In complex III, electrons pass from one cytochrome to the other cytochrome through an iron–sulfur protein. These electrons eventually reach cytochrome C. This process results in the transfer of protons from the mitochondrial matrix to the intermembrane space, which then generates electrochemical potential in the inner mitochondrial membrane through the free energy gained from the electron transfer. This electrochemical potential is used by mitochondrial ATP synthase to synthesize ATP (Nolfi-Donegan et al., 2020). The complex III consists of ten subunits in *S. cerevisiae*, including three catalytic subunits, COB (Cytochrome b), RIP1 (iron–sulfur protein), CYT1 (Cytochrome c1), and seven additional subunits: COR1, QCR2, QCR6, QCR7, QCR8, QCR9, and QCR10 (Brandt et al., 1994; Hunte et al., 2003). A previous study showed that complex III inhibition impaired the virulence and drug resistance of *C. albicans* in mice, and that the inhibition of cytochrome B significantly restricted the use of carbon sources, while the deletion of *RIP1* was found to have an effect on a range of virulence phenotypes of *C. albicans* (Vincent et al., 2016). In this study, we knocked out eight viable subunits, experimented with each mutant strain on the utilization of multiple alternative carbon sources and found that the *RIP1/COR1/QCR2/QCR7/QCR8* knockout strains all differed to varying degrees from the wild type in carbon-source utilization. However, a study of the effect of complex III on the virulence factor of *C. albicans* was found that QCR7 had a greater impact on biofilm formation and the maintenance hyphal growth on solid media than the other subunits. In addition, QCR7 is a protein that is directly and earliest interacts with fully hemimethylated cytochrome B in the assembly sites of the *S. cerevisiae* mitochondrial complex III and is involved in the conduction of protons from the matrix to the cytochrome b redox center (Lorusso et al., 1989). Although *C. albicans* is a significant pathogen, the function of *QCR7* in the virulence of *C. albicans* remains unknown.

This study, therefore, sought to explore the effects of QCR7 on the virulence of *C. albicans*,including its possible mechanism(s). We demonstrate the crucial involvement of *QCR7* in systemic infections due to *C. albicans*, in addition to its essential functions in the use of amino acids, N-acetylglucosamine (GlcNAc), and nonfermentable carbon sources. We used a *qcr7Δ/Δ* mutant that showed evident mitochondrial dysfunction. The *qcr7Δ/Δ* mutant exhibited significantly impaired biofilm formation and hyphal development. Furthermore, RNA-sequencing analysis identified several downregulated genes involved in carbohydrate transport and cell-surface functions, and that restore the corresponding pathogenic phenotype in case of the overexpression of the cell-surface-associated genes (*HWP1*, *YWP1*, *XOG1*, *SAP6*) in the *qcr7Δ/Δ* background. These data suggest that QCR7 regulates the cell-surface integrity to affect the use of carbon sources and pathogenic phenotypes, promoting host interaction.

## 2. Materials and Methods

### 2.1 Strains and growth conditions

Table S1 presents information on the strains, and Table S2 shows the primers used in this study. First, *C. albicans* mutants and complement strains were created following previous studies (Noble and Johnson, 2005; Nobile et al., 2009; Noble et al., 2010), and the overexpression strain was examined as previously described (Gerami-Nejad et al., 2013). In brief, the gene-knockout strategy of the present study involved gene knockout using polymerase chain reaction (PCR)-based homologous recombination. This strategy uses *C. albicans* SN152 with defective histidine (*HIS*), leucine (*LEU*), and arginine (*ARG*) genes as the parent strain and *HIS1 (Cd. HIS1*) from *C. dubliniensis, LEU2 (C. maltose LEU2, Cm.LEU2*) from *C. maltose*, and *ARG4* (*C. dubliniensis ARG4, Cd. ARG4*) from *C. dubliniensis* as the target genes to replace nutritional-marker genes for selection. This method typically knocks out the target genes one after the other, and two screening markers can be arbitrarily selected to knock out the target gene. In this study, the *LEU2* cassette from plasmid pSN40 was amplified, and the strain SN152 was transformed with the fusion PCR products of *LEU2* flanked by *QCR7* 5′ and 3′ fragments. The plasmid pSN52 was used as a PCR template to amplify the *HIS1* cassette, and the second *QCR7* allele was deleted using fusion PCR products of the *HIS1* marker flanked by *QCR7* 5′- and 3′-fragments (Figure S1). A copy of *QCR7* was returned to the *QCR7* locus to construct the gene-reconstituted strain. The entire *QCR7* open-reading frame, including 743 bp of the 5′end, was amplified. Next, the entire *QCR7* open-reading frame with upstream and the *ARG4* cassette from plasmid pSN69 and downstream flanks (approximately 400 bp) were amplified using fusion PCR assays. The PCR products were transformed into the *qcr7*Δ/Δ mutant to construct the *qcr7/QCR7* gene-reconstituted strain (Noble and Johnson, 2005) (Figure S2). The *qcr7Δ/Δ* mutant and reconstituted strain (*qcr7/QCR7*) were confirmed through PCR with the primer pairs shown in Supplementary Figure S3. However, the gene-overexpression strategy of the present study involved the amplification of the coding region of the corresponding genes by PCR and downstream cloning of the S. cerevisiae *ADH1* promoter to integrate inserts into a large intergenic region, NEUT5L which facilitates the integration and expression of ectopic genes (Gerami-Nejad et al., 2013) (Figure S4). For general growth and propagation, *C. albicans* strains were cultured at 30°C in yeast-extract–peptone–dextrose (YPD) medium (1% yeast extract, 2% peptone, 2% dextrose, and 2% agar).

### 2.2 Susceptibility assays

To evaluate the growth of *C. albicans* strains cultured with different carbon sources, strains were cultured overnight in YPD. Strains diluted to varying concentrations (5 μl of 10^7^ cells/ml to 10^3^ cells/ml) were spotted on yeast-extract–peptone solid medium (YEP; 1% yeast extract, 2% peptone). Next, washing and serial dilution with phosphate-buffered saline (PBS) were performed. This medium contained 2% glucose, 2%glycerol, 2% citrate, 2% acetate, 2% maltose, 2% ethanol, 2% lactate, and 50 mM GlcNAc. Subsequently, the sensitivity of *C. albicans* to cell-wall stress was assessed by spotting on YPD containing cell-wall stressors, such as Congo red (CR, 300 μg/ml), calcofluor white (CFW, 50 μg/ml), 0.04%SDS and caspofungin (0.25 μg/ml).

### 2.3 Biofilm formation assay

We evaluated the biofilm-formation mass of *C. albicans* using crystal-violet staining as described previously. First, strains were cultured overnight in liquid YPD at 30°C and 220 rpm and then harvested via centrifugation at 3,000 rpm after washing twice with sterile PBS. Subsequently, the mixture was resuspended in Spider liquid medium. Next, supplementation was performed using appropriate carbon sources and 2-mililiter *C. albicans*-strain suspensions (1 ml containing 5 × 10^6^ cells) and the suspensions were incubated overnight in 12-well flat-bottomed plates containing fetal bovine serum (FBS) before the final incubation for 90 min at 37°C (Silva, 2009; Silva et al., 2009; Abbes et al., 2017; Neji et al., 2017). Next, the nonattached cells were removed from the wells by washing once with 2 ml PBS, after which 2 ml of fresh corresponding induction medium was added to each well and incubated at 37°C for 48 h. Following the removal of medium, each well was washed once with PBS and treated with equivalent concentrations of methanol for 30 min. Next, equivalent concentrations of 1% crystal violet (Solarbio, China) were added to the wells for drying, and staining was performed for 1 h. Washing was performed under a gentle water flow until a colorless mixture was obtained, followed by incubation with equivalent amounts of acetic acid for 30 min to decolorize the mixture. Measurements were performed at 620 nm using a microplate reader.

### 2.4 Filamentous growth assays

The wild-type (WT) strain was cultured overnight in YPD, followed by washing and dilution with PBS to achieve a suspension with OD_600_ = 0.1. Next, the mixture was aliquoted in Spider liquid medium using appropriate carbon sources such as fermentable sugars (e.g., glucose, maltose, and sucrose) or alternative carbon sources (e.g., mannitol and GlcNAc) for hyphal induction at 37°C. The cells were harvested, and hyphal morphologies were visualized under a fluorescence microscope (Olympus, Japan) at indicated time points. Subsequently, each cell type (1 × 10^5^ cells in 5 μl of PBS) was spotted on Spider solid medium with appropriate carbon sources. The cultures were then incubated at 37°C, and photographs were taken after 7 days.

### 2.5 Measurement of mitochondrial membrane potential

To determine the mitochondrial membrane potential (MMP), the MMP assay kit with JC-1 (Beyotime Institute of Biotechnology) was used. MMP decreases can be easily detected through JC-1’s transition from red to green fluorescence. Cells were collected when strains subcultured in fresh medium reached the log phase of growth and incubated with equal volumes of JC-1 dyeing solution at 37°C for 15 min in the dark. Cells were washed twice and resuspended in JC-1 dyeing solution. Next, we explored the emission spectra at 488 nm and excitation spectra at 595 nm using the Cytomics FC500 flow cytometer (Beckman Coulter); the red/green mean fluorescence intensities (FIs) were recorded for each sample.

### 2.6 Measurement of intracellular-reactive-oxygen-species levels

All strains were cultured overnight in YPD at 30°C and 220 rpm, followed by washing and resuspension in PBS. Subsequently, 2 × 10^6^ cells were stained using fluorescent dichlorodihydrofluorescein diacetate dye (20 μg/ml). Next, these cells were incubated at 37°C for 20 min in the dark. Subsequently, FIs were recorded using a multifunctional enzyme-mark instrument (SpectraMax Paradigm) with an excitation wavelength of 480 nm and emission wavelength of 530 nm. Finally, the levels of reactive oxygen species (ROS) were determined by evaluating each strain thrice.

### 2.7 Measurement of intracellular ATP contents

All strains were cultured overnight in YPD at 30°C. Cells were then subcultured in fresh medium until the log phase of growth was reached. Next, 2 × 10^6^ cells of each strain were mixed completely with equal volumes of the BacTiter-Glo reagent (Promega Corporation, Madison, WI), followed by incubation at room temperature for 15 min in the dark. Finally, luminescent signals were detected using the full-wavelength multifunctional enzyme-mark instrument (SpectraMax Paradigm).

### 2.8 Sap-activity-testing assays

Secreted aspartyl protease (Sap) activity assays were performed using 0.17% yeast-nitrogen-base (YNB) medium (without aa or AS) + 0.1% BSA, containing 2% glucose as the carbon source and bovine serum albumin (BSA])as the sole nitrogen source (Crandall and Edwards, 1987). Water was used to suspend 1 × 10^5^ cells, after which they were spotted on YNB–BSA agar plates and incubated at 37°C for 5 days. The size of the white halo rings indicated the activity level. This experiment was repeated thrice.

### 2.9 RNA isolation, cDNA library preparation, and sequencing

Through transcriptome sequencing, we evaluated the global transcriptional response of *C. albicans* in RPMI 1640. All strains were cultured overnight in YPD at 30°C and subcultured in fresh YPD (buffered) at 30°C. Subsequently, *C. albicans* cells at the log phase of growth were resuspended in RPMI 1640, followed by incubation at 37°C for 4h according to the manufacturer’s instructions. Next, RNA extraction was performed using TRIzol (TIANGEN). In brief, 5 × 10^6^ cells were isolated and added to a precooled mortar. The cells were then grounded to powder with liquid nitrogen, after which the sample was stirred in 1 ml TRIzol for 10 min. Subsequently, chloroform (200 μl) was added to separate the aqueous RNA-containing solution from the intermediate and organic layers, which was followed by RNA precipitation using isopropyl alcohol (equal volume) and the addition of 1 ml ethanol (75%) to each RNA pellet. Finally, the samples were centrifuged (12,000 rpm/5 min/4°C), and the RNA was resuspended in 20 μL nuclease-free water and then stored at −80°C. Furthermore, the RNA samples were assessed using Agilent 2100 Bioanalyzer (Agilent Technologies, Santa Clara, CA, USA) and NanoPhotometer spectrophotometer (Implen). Triplicate samples used in all assays were used to construct an independent library, and the sequencing and analysis were performed. The libraries for sequencing were constructed using the NEB Next Ultra RNA Library Prep Kit for Illumina (NEB). The poly (A)-tailed mRNA was enriched using the NEB Next Poly (A) mRNA Magnetic Isolation Module kit. The mRNA was fragmented into segments of nearly 200 bp. First-strand cDNA was synthesized from the mRNA fragments through reverse transcription using random hexamer primers. Next, second-strand cDNA was synthesized using DNA polymerase I and RNaseH. The end of the cDNA fragment was subjected to an end repair that included adding a single “A” base, followed by adapter ligation. PCR was used to purify and enrich the products to construct a DNA library. The final libraries were quantified using Qubit 2.0 Fluorometer (Invitrogen) and Agilent 2100 Bioanalyzer. After performing validation through quantitative reverse transcription-PCR, libraries were subjected to paired-end sequencing with a reading length of 150 bp using Illumina Novaseq 6000 (Illumina).

### 2.10 Quantitative real-time PCR

All the strains were cultured overnight in YPD at 30°C and subcultured in fresh YPD (buffered) at 30°C until growth at the log phase. Subsequently, cells were resuspended in RPMI 1640, followed by incubation at 37°C for 4h. The methods for RNA extraction were those described above. Extracted RNA was treated with DNase I (Takara) to remove contaminating DNA, and cDNA was synthesized with a PrimeScript™ RT Reagent Kit (Takara) following the manufacturer’s protocol. The cDNA was used as template for RT–qPCR using a SYBR Green mix kit (Mei5 Biotechnology Co., Ltd.) on a CFX Connect Real-Time PCR Detection System. The 18S rRNA genes were used as endogenous controls as specified. The results were analyzed using the 2^-ΔΔCT^ method. The primers for the genes used in the RT-qPCR are shown in Supplementary Table S2.

### 2.11 Statistical analysis

All experiments were biologically replicated thrice. Data were presented as means + standard errors of the means. Statistical analyses were performed using GraphPad Prism 8 software. Significant differences were calculated using two-way analyses of variance and Student’s *t*-tests.

## 3. Results

### 3.1 Complex III plays an important role in the virulence factors of *Candida albicans*

Previous studies identified that disabling Complex III can prevent fungal adaptation to nutrient deprivation, sensitize *C. albicans* to attack by macrophages, and curtail virulence in mice (Vincent et al., 2016). However, in these studies, the catalytic subunit RIP1 was chosen as a representative and used in the remaining experiments. It is unclear whether other subunits are equally important for the functional performance of mitochondrial Complex III (CIII) and the maintenance of *C. albicans* virulence. In this study, we identified ten genes (*COB, RIP1, CYR1, COR1, QCR2, QCR6, QCR7, QCR8, QCR9*, and *QCR10*) encoding the subunit of mitochondrial Complex III using the Candida Genome Database (CGD, http://www.candidagenome.org/). Among them, apart from *cobΔ/Δ* and *cyr1Δ/Δ* mutants, which were inviable, we successfully knocked out the other eight subunits and tested the utilization of several carbon sources as energy sources for the pathogens. As shown in Figure 1A, in the right panel, in media with glucose, sucrose, and mannitol as carbon sources, *rip1Δ/Δ, cor1Δ/Δ, qcr2Δ/Δ, qcr7Δ/Δ*. and *qcr8Δ/Δ* showed a similar lag phase. However, the above mutant displayed remarkable growth defects compared to that of the WT in media with other host-relevant carbohydrates (e.g., maltose, citrate, acetate) as carbon sources. To examine the effect of Complex III mutants on the major virulence factors associated with *C. albicans*, the panel of mutants was examined for their biofilm-formation ability, hyphal growth, etc. Consistent with an alteration in carbon assimilation, the corresponding mutants also displayed defects in hyphal growth on solid media and were unable to form the dense matrix of the biofilm (Figure 1B-C). The remaining subunit formed Complex III; these coding gene mutants showed a similar phenotype to the wild-type strain. Surprisingly, we found that the absence of *QCR7* had a more evident phenotypic impact on the ability of *C. albicans* to maintain hyphal growth on solid media and biofilm formation than the other subunit knockout strains. The results indicate that QCR7 is critical to virulence in *C. albicans*.Thus, a series of experimental studies were conducted on *QCR7*, as described below.

**Figure 1.**
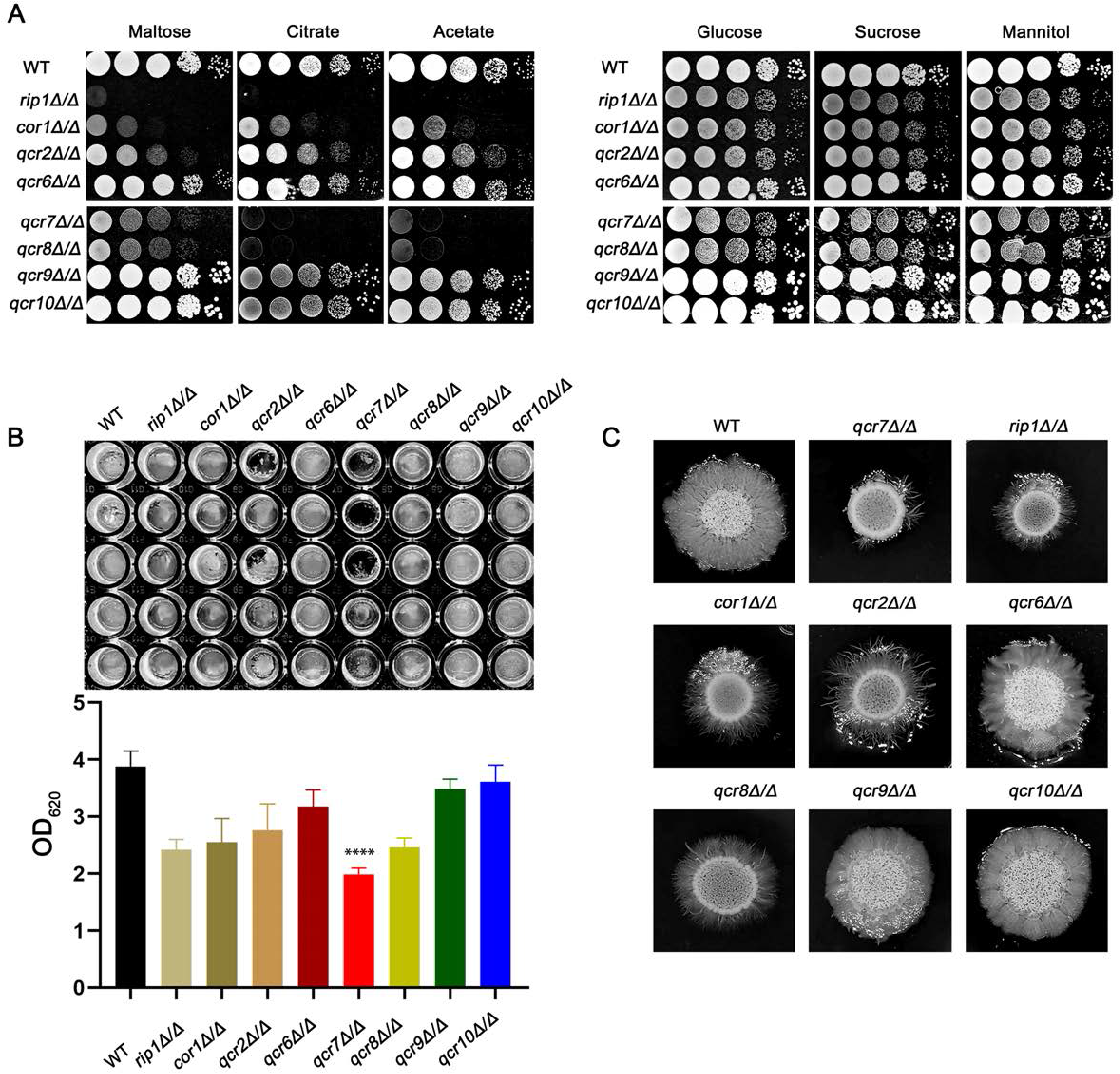
Complex III plays an important role in the virulence of *Candida albicans*. (A) Strains were cultured overnight in YPD, followed by washing and serial dilution with PBS. Different *rip1Δ/Δ, cor1Δ/Δ, qcr2Δ/Δ, qcr6Δ/Δ, qcr7Δ/Δ, qcr8Δ/Δ, qcr9Δ/Δ*, and *qcr10Δ/Δ* concentrations (5 μl of 10^7^ cells/ml to 10^3^ cells/ml) were spotted on solid YP media containing 2% glucose and several alternative carbon sources. Plates were then photographed after incubation at 30°C for 2 days. (B) *C. albicans* suspension of 5 ×10^6^ cells in Spider medium were incubated in 12-well flat-bottom plates at 37°C for 90 min, after which nonattached cells were removed from the wells by washing once with PBS. Fresh corresponding induction medium was added to each well and incubated at 37°C for 48h. Next, the well was washed with PBS and stained with crystal violet; after decolorization, measurements were taken using a microplate reader at 620 nm. “*” represents *p* < 0.05 and “****” represents *p* < 0.0001 for the WT vs. mutant strains. (C) Cells of each type (1 × 10^5^ cells in 5 μl PBS) were spotted on regular indicated filament-inducing agar plates and incubated at 37°C for 7 days before taking photographs.

### 3.2 *QCR7* mutants attenuated the virulence of *C. albicans* infection in vivo

Based on the above findings, it is reasonable to assume that *QCR7* mutant inhibits fatal *C. albicans* infections. The inhibition of Complex III was found to be effective in attenuating the virulence of *C. albicans* in mice in previous reports (Vincent et al., 2016). Therefore, we performed relevant experiments and analyzed in a mouse model of systemic candidiasis. The survival rate of the mice injected with the *qcr7Δ/Δ* mutant was much higher than that of those injected with the WT and *QCR7* complement strains. Furthermore, no mice died within 3 weeks of infection with the *qcr7Δ/Δ* mutant (Figure 2A). During pathological characterization, the kidneys, the main target organ, of the infected mice served as the key indicators of host infection (Leunk and Moon, 1979). Regarding the kidney anatomy, the mice injected with the *qcr7Δ/Δ* mutant had smaller kidneys than those injected with the WT strain (Figure 2B). To quantitatively evaluate the invasive effects of *C. albicans* on the kidney, we investigated the fungal loads that correlated with the severity of kidney failure. By calculating the total number of colony-forming units (CFU) in the kidneys on day 2, we observed that the fungal burden in the kidneys injected with the *qcr7Δ/Δ* mutant significantly reduced (Figure 2C-E). A histological analysis of the kidney-tissue samples obtained from the mice infected with the WT strain also revealed large necrosis and areas with inflammatory cell infiltration (Figure 2F).

**Figure 2.**
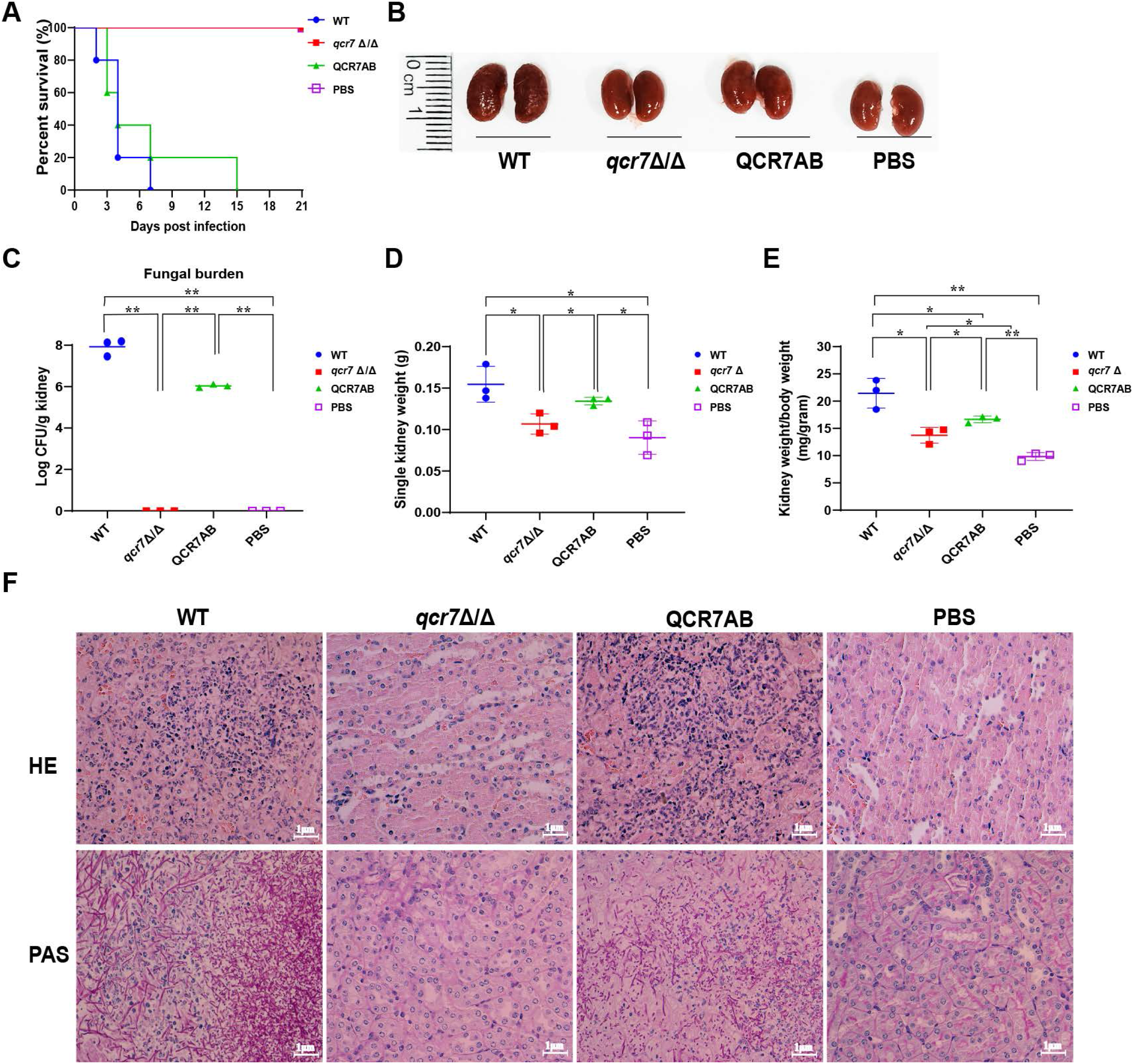
*QCR7* mutant attenuates the virulence of *C. albicans* infection in vivo. (A) Mouse survival following illness initiation with the injection of 5 × 10^5^ *C. albicans* cells. (B) Comparison of mouse kidneys was conducted by injection with WT, *qcr7Δ/Δ*, and QCR7AB. Mice were sacrificed 48h after the *C. albicans* infection. (C-E) Fungal burden in the kidneys of mice injected with WT, *qcr7Δ/Δ*, and QCR7AB. The mice were sacrificed 2 days after an intravenous injection with 5 × 10^5^ CFU *C. albicans*. Kidney weights were calculated as each mouse’s weight per gram of body weight. “*” represents *p* < 0.05 and “**” represents *p* < 0.01 for the WT vs. mutant strains. (F) Representative hematoxylin–eosin- or periodic acid-Schiff-base-stained sections of kidney-tissue samples from three experiments.

The kidney tissues obtained from the *qcr7Δ/Δ* mutant-infected mice showed significantly less necrosis and tissue damage. Periodic acid-Schiff-base staining consistently revealed more necrotic areas in the kidney, which differed between the WT-strain-infected mice with several *C. albicans* hyphae, pseudo-hyphae, and spores, and the *qcr7Δ/Δ* mutant-infected mice (Figure 2F). Therefore, these results suggest that virulence reduction of the *qcr7Δ/Δ* mutant results from defective tissue invasion and colonization ability.

### 3.3 *QCR7* deletion affected the mitochondrial function and carbon-source-usage pattern of *C. albicans*

Fungi can use carbon sources to obtain energy for anabolic metabolism through glycolysis and aerobic respiration. The energy generated through the electronic transport chain is the primary source of energy for fungal growth. Therefore, if mitochondrial dysfunction affects aerobic respiration, the glycolytic pathway becomes the main metabolic pathway for energy metabolism. In addition, glycolysis is crucial for carbon assimilation, and this pathway also influences the invasiveness and virulence of *C. albicans* (Barelle et al., 2006). *C. albicans* uses multiple carbon sources in vitro, including glucose, GlcNAc, carboxylic acids (e.g., lactate), and amino acids (Williams and Lorenz, 2020). Compared with the WT strain, the *qcr7Δ/Δ* mutant grew slightly slower in the glucose-, sucrose-, and mannitol-only media. However, the growth of the mutant in the media containing GlcNAc or other non-glucose carbon sources was more significant than that of the WT strain (Figure 3A). The *qcr7Δ/Δ* mutant did not use amino acids as carbon sources (Figure S5). The results indicate that the *qcr7Δ/Δ* mutant has a defective respiratory metabolism, which may affect ATP production during carbon metabolism. Furthermore, this implies that *QCR7* deletion does not substantially regulate the in vitro growth of *C. albicans* in the presence of alternative carbon sources.

**Figure 3.**
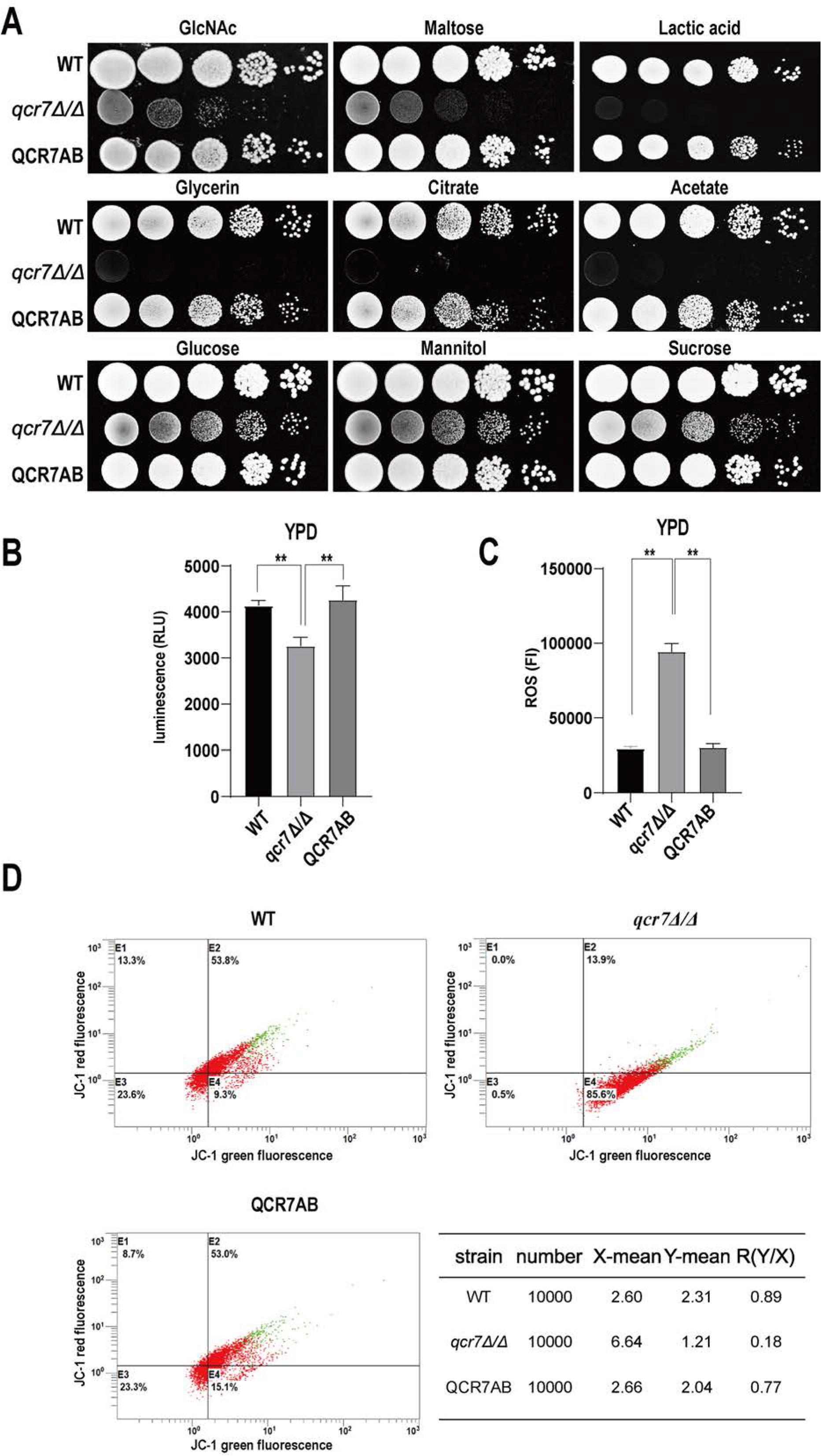
*QCR7* is required for maintaining mitochondrial functions and carbon-source use. (A) Strains were cultured overnight in YPD, followed by washing and serial dilution with PBS. Different WT, *qcr7Δ/Δ*, and QCR7AB concentrations (5 μl of 10^7^ cells/ml to 10^3^ cells/ml) were spotted on solid YP media containing 2% glucose and several alternative carbon sources. Plates were then photographed after incubation at 30°C for 2 days. (B) The intracellular ATP content was measured using a microplate reader. “**” represents *p* < 0.01 and “****” represents *p* < 0.0001 for the WT vs. mutant strains. (C) ROS levels were measured using the dichlorodihydrofluorescein diacetate dye and a multifunctional enzyme-mark instrument. (D) Mitochondrial membrane potential (MMP) was evaluated using the JC-1 assay kit and Cytomics FC500 flow cytometer (Ex/Em of 595/488 nm), and the red/green mean fluorescent intensities were recorded for each sample. The transition of JC-1 dye from red to green fluorescence was used to easily detect the decrease in MMP, after which MMP was determined as the ratio of red to green fluorescence.

Because QCR7 is a component of the mitochondrial respiratory chain, we examined the intracellular ATP content, endogenous ROS production, and mitochondrial membrane potential (MMP) of *qcr7Δ/Δ* mutants to determine their effect on mitochondrial functions. As expected, a significant decrease in intracellular ATP contents was observed in the *qcr7Δ/Δ* mutant (Figure 3B). Moreover, after evaluating the MMP using the JC-1 assay kit, we observed that the MMP of the mutant was significantly decreased compared with that of the WT strain (Figure 3D). In addition, the *qcr7Δ/Δ* mutant produced a higher level of ROS (Figure 3C). These data suggest the important function of QCR7 in maintaining mitochondrial homeostasis.

### 3.4 *QCR7* deletion resulted in impaired biofilm formation and hyphal-growth maintenance after the exposure of *C. albicans* to various carbon sources

The biofilms of *C. albicans* are highly structured; they contain yeast forms of cells, pseudohyphal cells, and hyphal cells surrounded by an extracellular matrix (Fox and Nobile, 2012). Furthermore, *C. albicans* biofilms functions as a reservoir of drug-resistant cells that can isolate, multiply, and cause bloodstream infections (Lohse et al., 2018). In addition to forming biofilms on implanted medical devices, the surface of the host is also a major location for the formation of *C. albicans* biofilms. The transition from commensal to pathogenic *C. albicans* requires effective adaptation to the host environment. It has been observed that carbon plays a central role in metabolism and is key to host-surface colonization during infection (Brown et al., 2014). Glucose and other sugars influence the ability of *C. albicans* to form biofilms (Ng et al., 2016; Pemmaraju et al., 2016). Moreover, the biofilm’s thickness and structure vary with different carbon sources (Pemmaraju et al., 2016). Thus, to understand the importance of *QCR7* in biofilm formation, we performed an additional biofilm analysis under multiple conditions. Defective biofilm formation was evident in the *qcr7Δ/Δ* mutant when incubated in the Spider medium supplemented with glucose, mannitol, GlcNAc, maltose, and sucrose (Figure 4A). Therefore, these results indicate the importance of *QCR7* in appropriate carbon-induced biofilm formation by *C. albicans*.

**Figure 4.**
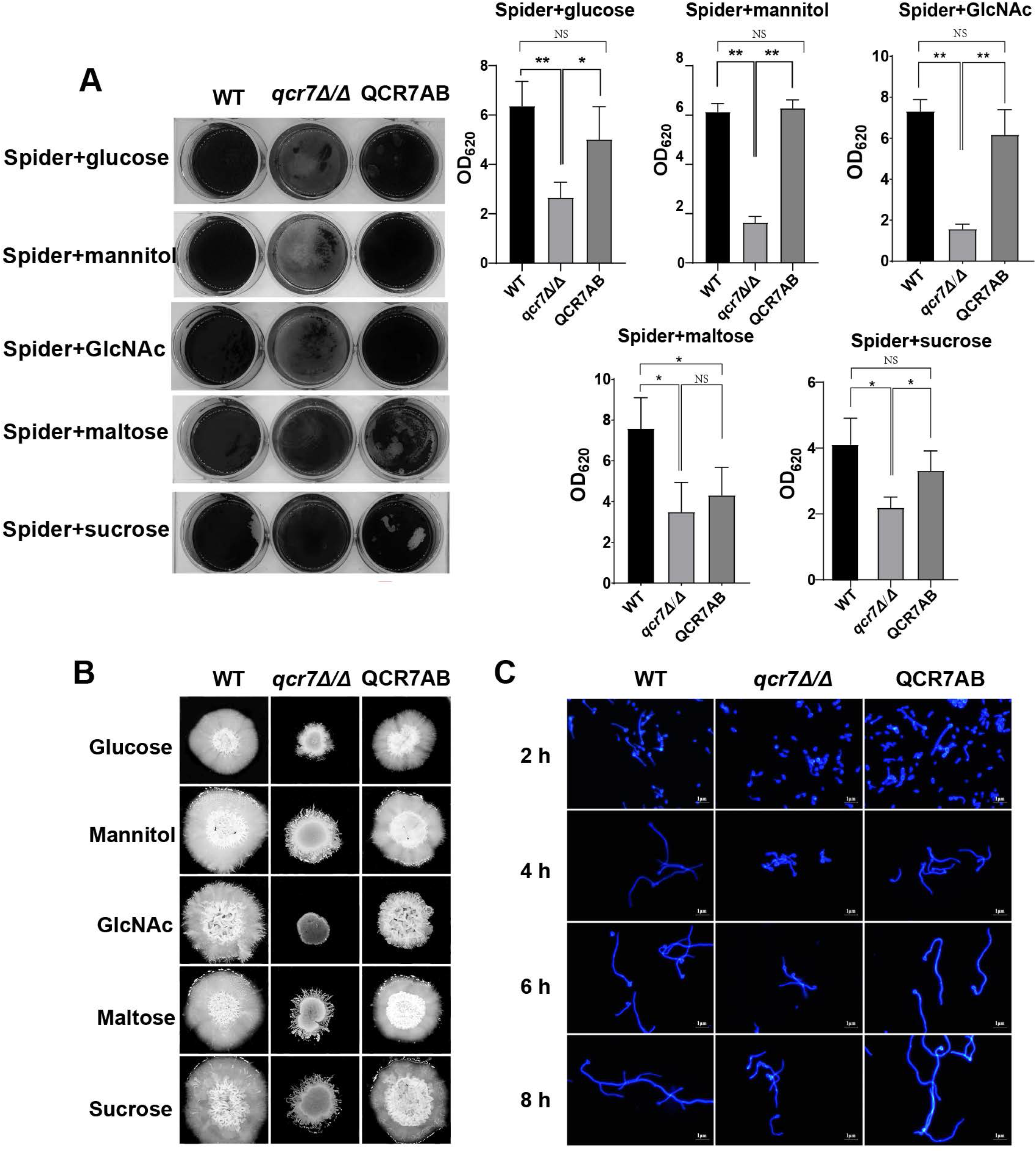
Deletion of QCR7 affects biofilm formation and hyphal growth maintenance in Candida albicans under various carbon sources, with the most significant effect under GlcNAc conditions. (A) *C. albicans* suspensions of 5 × 10^6^ cells in Spider medium supplemented with mannitol, glucose, sucrose, maltose, and GlcNAc as sole carbon sources were incubated in 12-well flat-bottom plates at 37°C for 90 min, after which nonattached cells were removed from the wells by washing once with PBS. Fresh corresponding induction medium was added to each well and incubated at 37°C for 48h. Next, each well was washed with PBS and stained with crystal violet; after decolorization, measurements were taken using a microplate reader at 620 nm. “*” represents *p* < 0.05 and “**” represents *p* < 0.01 for the WT *vs*. mutant strains. (B) Each cell type (1 × 10^5^ cells in 5 μl of PBS) was spotted on the Spider medium, containing 2% glucose, 2% mannitol, 2% maltose, 2% sucrose, and 2% GlcNAc at 37°C. Photographs were then taken after 7 days of growth. (C) The wild type, *qcr7Δ/Δ*, and QCR7AB were cultured overnight in liquid YPD at 30°C, followed by washing and dilution to OD_600 nm_ = 0.1 of PBS, after which resuspension in Spider medium supplemented with GlcNAc as the sole carbon source was conducted, and incubation continued at 37°C. The hyphal morphology was finally visualized through fluorescence microscope (Olympus, Japan). Scale bars are 1 μm.

*C. albicans* can form hyphal cells both in planktonic cultures and during the maturation step of biofilm formation (Sudbery, 2011). As reported previously, the yeast accounting for the hyphal transformation of *C. albicans* is key to causing invasive infections by invading epithelial cells and causing tissue damage. Therefore, the ability of fungi to form hyphae is evaluated through their filamentous growth abilities in liquid cultures or colonial morphology on solid media (Sudbery, 2011). In this study, we assessed the influence of *QCR7* on the hyphal induction of *C. albicans* cultured in media containing various carbon sources. As expected, the results showed that the differences in hyphal morphology were more evident on the solid media. After culturing at 37°C, although the WT formed significant wrinkled colonies and hyphal extensions, the mutant formed only smooth colonies and slight hyphae.Particularly under GlcNAc, the colony derived from the *qcr7Δ/Δ* mutant was smooth (Figure 4B). Accordingly, the *qcr7Δ/Δ* mutant showed shorter and sparser hyphae when incubated in a liquid-induced medium with GlcNAc as the sole source of carbon (Figure 4C). However, no significant differences were observed in the hyphal length of the fungi cultured in liquid media with other carbon sources (Figure S6) compared with the colony formed by the *qcr7Δ/Δ* mutant, where the WT strain developed wrinkled macro-colonies and extended the hyphae on solid media more evidently. In addition, this was accompanied by the loss of hyphal-related hydrolase activity (Figure S7). Therefore, the *qcr7Δ/Δ* mutant might have affected the hyphal growth because of its impaired use of carbon sources.

### 3.5 Comparative transcriptome analysis provided information on the *QCR7*-associated virulence mechanisms of *C. albicans*

The transcriptome of the *qcr7Δ/Δ* mutant was compared with that of the WT strain in media mimicking the in vivo conditions. The results indicated that 307 genes were downregulated and 54 genes were upregulated with expression-fold changes of >2.0 and <−2.0 (P ≤ 0.05), respectively, in the *qcr7Δ/Δ* mutant compared with the WT strain. Therefore, we subjected these gene sets to pathway analysis using Gene Ontology (GO). Among the 307 downregulated genes, the most enriched GO term was transport (22.5%). The enriched terms also included processes such as translation (21.8%), ribosome biogenesis (20.5%), RNA metabolic processes (20.2%), the regulation of biological processes (18.9%), organelle organization (15.6%), response to stress (13.0%), and lipid metabolic processes (12.1%), followed by those related to filamentous growth (10.4%), cellular protein modification processes, response to chemicals, cellular homeostasis, biofilm formation (4.6%), carbohydrate metabolic processes (4.2%), and cell adhesion (2%) (Figure 5A). Furthermore, the list of the 54 upregulated genes list comprised 26 genes with unknown functions. The eight highest gene categories that were downregulated included those related to transport (33.3%), the regulation of biological processes (22.2%), cellular homeostasis (11.1%), protein catabolic process (9.3%), and responses to chemicals, including those related to filamentous growth, carbohydrate metabolic process, and cell adhesion (1.9%) (Figure 5B). An analysis of the GO category “cellular components” also found GO terms associated with the membrane and cell wall to be significantly (P ≤ 0.05) enriched in the downregulated gene sets (Figure S8). Regarding the process related to sugar transportation through the plasma membrane, it can be proposed that membrane downregulation by some genes can affect the use of sugar. Previous evidence showed that mitochondrial function directly influences the cell wall (Duvenage et al., 2019). In addition, the absence of the Complex I regulator Goa1 (She et al., 2016) or the mitochondrial sorting and assembly machinery subunit Sam37 (Qu et al., 2012) results in defective cell-wall integrity (CWI), suggesting that the mitochondrial gene defects affecting the cell wall also apply to the Complex III regulator QCR7. Intriguingly, an analysis of the downregulated gene sets related to filamentous growth and biofilm formation revealed that significantly downregulated genes were involved in regulating biofilm formation (*SAP6, HWP1, XOG1, HYR1, YWP1, ALS3, CSH1, PGA10*, and *SUN41*) and filamentous growth (*ENO1, SSB1, DDR48, ALS3*, and *SUN41*); these genes were also identified as cell-wall-related genes, indicating that impaired filamentous growth and biofilm formation were partly affected by the downregulation of cell-wall-associated genes. Moreover, several genes encoding glycosylphosphatidylinositol-anchored cell-wall proteins that influenced the cross-linking of cell-wall components were also downregulated (*PGA3, PGA6, PGA7, PGA10, PGA34, PGA35, PGA53, PGA54, PGA56, PGA59*, and *PGA63*) (Figure S8). To verify whether QCR7 deletion negatively affects CWI, we spotted it on a medium containing cell-wall stressors, such as CFW, CR, caspofungin, and SDS. We observed that disrupting QCR7 led to damaged cell-wall biosynthesis (Figure S8).

**Figure 5.**
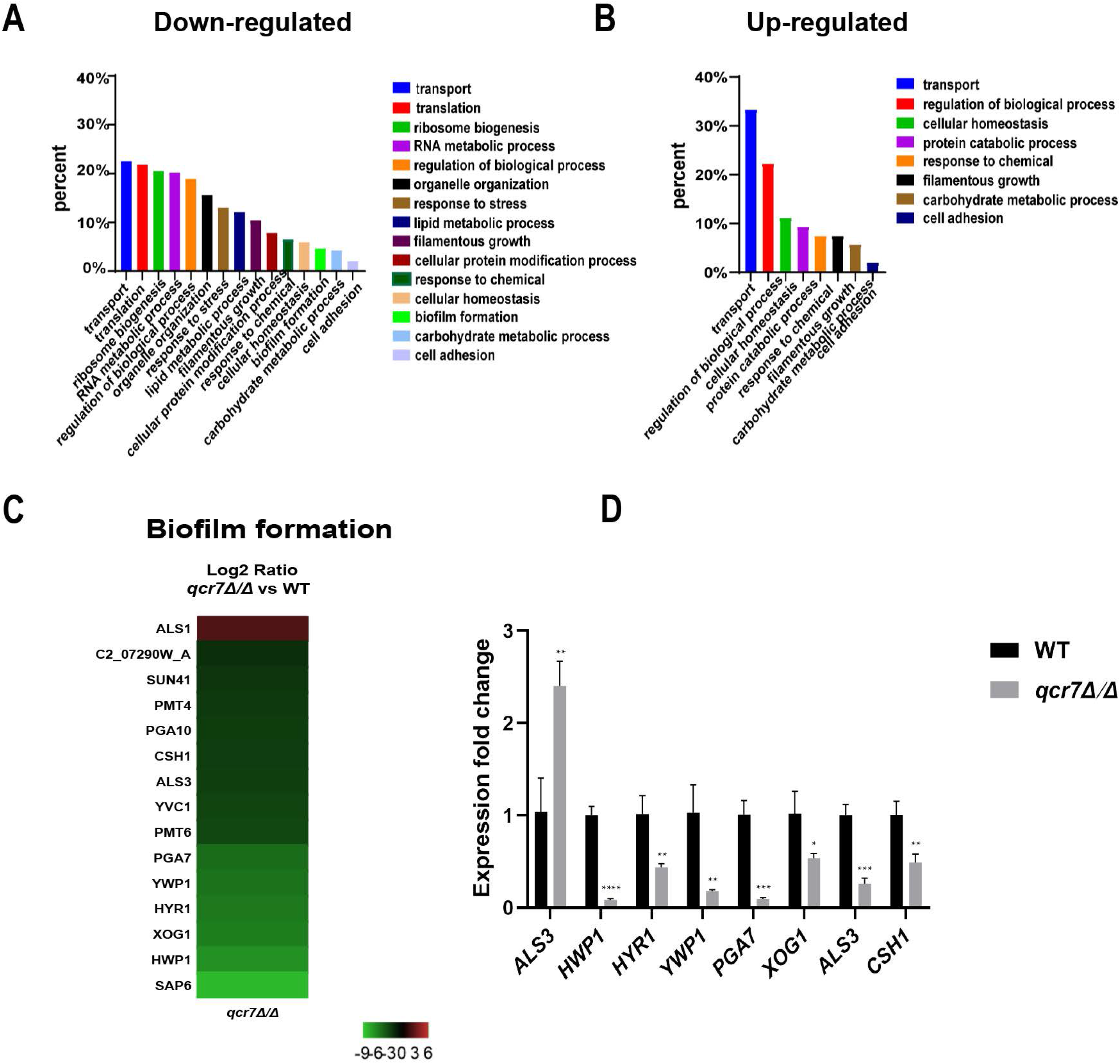
Comparative transcriptomic analysis of *QCR7* mutants of *C. albicans*. The distribution of significantly downregulated (A) or upregulated (B) genes in the GO category “molecular process” following *qcr7Δ/Δ*, compared with the wild type. (C) A heatmap displaying the upregulated and downregulated genes that associate partly with the GO category “biofilm formation”. Fold enrichment of these genes is shown in Table S3. (D) Validation of the genome-wide transcriptional data by qPCR in the *qcr7Δ/Δ* mutant strain.

Furthermore, our RNA-sequencing data revealed the most significant downregulation of several genes of biofilm formation, and we confirmed the reliability of the results by RT-qPCR (Figure 5C). In *C. albicans*, biofilm development requires six master transcriptional regulators (BCR1, BRG1, NDT80, ROB1, TEC1, EFG1) (Nobile et al., 2012). We further investigated whether six core biofilm regulatory factors in *C. albicans* regulate QCR7 expression. As shown in Figure S9, the deletion of these master regulator significantly affects *QCR7* expression. In addition, we overexpressed *QCR7* in these transcription-factor-encoding knockout strains and performed biofilm formation assays, showing that *QCR7* could partially restore the biofilm deficiency caused by *NDT80* deletion (Figure S10). These data demonstrated the role of *QCR7* in biofilm formation.

### 3.6 QCR7-regulated genes in cell surfaces play a significant role in biofilm formation, hyphal growth on solids, and the use of carbon sources

To further identify downstream biofilm-formation-related genes regulated by *QCR7* and the probable mechanisms involved, we selected five genes (*HWP1, YWP1, XOG1, HYR1*, and *SAP6*) known to play an important role in biofilm formation and that were downregulated with an expression-fold change of < −5.0 (P≤0.05) in the *qcr7Δ/Δ* mutant compared with the WT strain based on the transcriptome and overexpression of theses gene in the *qcr7Δ/Δ* background. We found that the overexpression of four genes (*HWP1, YWP1, XOG1*, and *SAP6*) partly but significantly rescued the *qcr7Δ/Δ* biofilm phenotype, but that *HYR1* overexpression in this genetic background suppressed the biofilm phenotype of the *qcr7Δ/Δ* mutant even more (Figure 6A). Furthermore, the partial restoration of the phenotype also occurred in hyphal growth in solid medium. The increased expression levels of *HWP1, YWP1, XOG1*, and *SAP6* in the *qcr7Δ/Δ* mutant formed a halo of more abundant and longer agar-invading hyphae around the colony center than those seen with the *qcr7Δ/Δ* mutant, whereas the overexpression of *HYR1* showed no formation of a halo of agar-invading hyphae (Figure 6B). Considering whether the same mechanism regulates the reduction of each virulence phenotype, and restricted carbon source utilization, we tested for growth on several alternative carbon sources to examine the utilization of the carbon sources by these strains. The results showed that carbon use was consistent with the virulence phenotype, with four genes (*HWP1, YWP1, XOG1, SAP6*) overexpressed in the *qcr7Δ/Δ* background showing a partial reversal of the use of these carbon sources, while the overexpression of *HYR1* inhibited the use of the carbon source even more (Figure 6C). The mechanism behind this is not yet clear.

**Figure 6.**
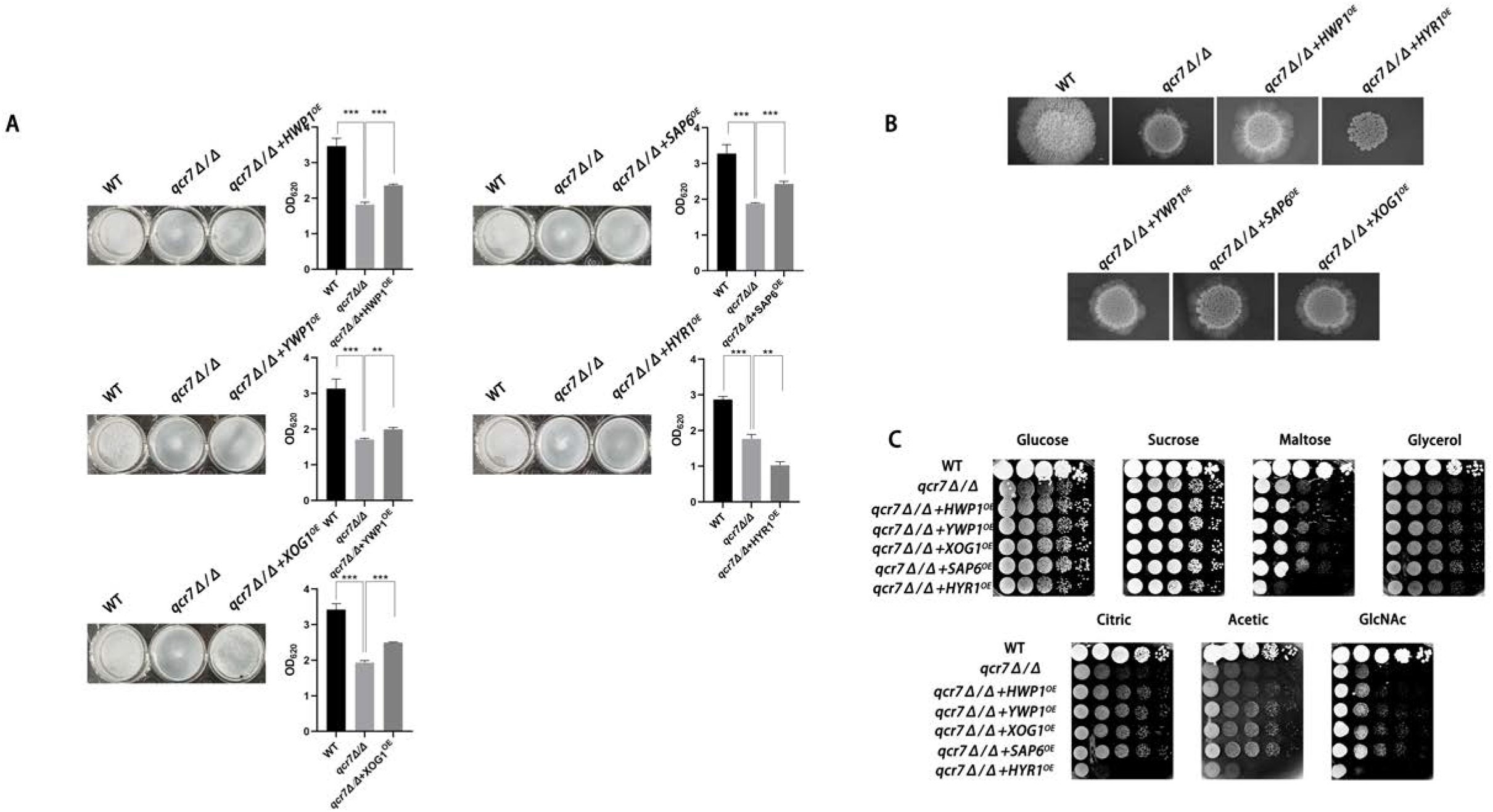
Biofilm formation, hyphal growth on solid, and spot assay on cell-wall stressors or multiple alternative carbon sources with WT, *qcr7Δ/Δ, HWP1, YWP1, XOG1, SAP6*, and *HYR1* overexpressing strains in the *qcr7Δ/Δ* background. (A) Biofilm formation (48h, 37°C) of strains was profiled and statistically compared to reference strain *C. albicans* SN250. (B) Each cell type (1 × 10^5^ cells in 5 μl of PBS) was spotted on the Spider medium at 37°C. Photographs were then taken after 7 days of growth. (C) The sensitivities of wild-type strain, *qcr7Δ/Δ* strain, and overexpression strain to different carbon-source conditions were observed by spot assay and incubation was conducted at 30°C for 2 days before photographs were taken.

Altogether, the above results confirm that the restricted carbon utilization caused by the deletion of *QCR7* is related to the impairment of the virulence phenotype and cell-surface changes.

## 4. Discussion

In this study, we constructed eight subunit-encoding genes mutant that are viable in Complex III and investigated the effects of the deletion of these genes on the ability of carbon-source utilization and on key virulence phenotypes of *C. albicans*. We found that the deletion of RIP1, a catalytic subunit of Complex III, had a significant effect on both the carbon-source utilization and virulence of *C. albicans*, which was consistent with a previous study (Vincent et al., 2016). While we were surprised to find that the deletion of QCR7, one of the seven non-catalytic subunits of Complex III, had a phenotypic deficiency in the ability of *C. albicans* to maintain hyphal growth on solid media and biofilm formation compared to the other subunit knockout strains, we first considered this in terms of the constitutive structure of QCR7 as a direct effect. QCR7, the earliest protein to interact with the fully hemylated Cytb (Hildenbeutel et al., 2014), is present in all bc1 Complex assembly intermediates containing nuclear coding subunits and therefore plays an important role in the assembly of Complex III. In order to gain a deeper understanding of the impact of the QCR7 knockout on the biological and pathogenic functions of *C. albicans*, we then carried out a series of studies. Because the preliminary results based on evident phenotypic changes, which relate to virulence factors, suggested that QCR7 deletion reduces the invasive infection of C. albicans, we used mice as the host systems to assess the effects of QCR7 deletion on the virulence of C. albicans in vivo. Although all the mice infected with the mutant survived beyond 3 weeks after infection, those infected with the WT strain died within a week. A histological analysis and fungal-kidney-tissue loads revealed the lack of Candida cells in the kidneys of the mice infected with the *qcr7Δ/Δ* mutant, showing significantly less necrosis and less-severe tissue damage. Therefore, the ability of the mutants to invade and colonize the host tissues might have been weakened.

Carbon sources are extremely diverse in different host environments, making it important for *C. albicans* to use alternative carbon sources. Although it inhabits niches with limited glucose availability, it is also found in environments rich in alternative carbon sources. Therefore, this fungus uses nonfermentable carbon sources, such as acetate or lactate, to adapt to host environments (Piekarska et al., 2006). Alternatively, the efficient assimilation of multiple nonfermentable carbon sources enhances its virulence (Ramírez and Lorenz, 2007). GlcNAc, an alternative carbon source, can be easily found in the mucosal membranes of human hosts (Sengupta and Datta, 2003), and it can strongly induce hyphal morphogenesis (Williams and Lorenz, 2020). In addition, studies have reported that *C. albicans* can use amino acids as carbon sources, which is crucial for its virulence (Vylkova et al., 2011). To further determine the effect of QCR7 deletion on carbon-source utilization, a wild-type allele was restored to the QCR7 knockout strain and tested in a more diverse-carbon-source medium. It was found that QCR7 complementation can compensate for QCR7 deficiency, resulting in growth-lag phase on glucose, mannitol, and sucrose as carbon sources and growth defects on non-fermentable carbon sources (glycerin, citrate, and acetate), GlcNAc, lactate and amino acids as carbon sources. The reduced ability to utilize carbon sources raised the question of whether the energy supply within the strain is affected by the deletion of QCR7. The mitochondria are the primary sources of energy for cell growth, apoptosis, proliferation, and other cellular processes. Therefore, mitochondrial function is crucial in regulating the morphology and virulence of *C. albicans*. The assay of mitochondrial function suggested that QCR7 deletion affected intracellular ATP content, which may be related to QCR7 as a subunit in the electron transport chain, affecting the level of mitochondrial membrane potential (MMP), and the elevated intracellular ROS levels also further confirmed the dysfunctions in the mitochondria.

We further observed that the mitochondrial dysfunction induced by the QCR7 knockout impaired the biofilm formation of the *C. albicans*. Compared with the WT strain, the QCR7 deletion diminished biofilm formation in the Spider medium supplemented with mannitol, glucose, GlcNAc, maltose, and sucrose as the sole carbon source. Robust hyphal formation is necessary for *C. albicans*-biofilm maturation. Furthermore, the *qcr7Δ/Δ* mutant showed significantly different hyphal morphologies when stimulated by multiple alternative carbons. Similar phenotypic differences in biofilm formation were also observed in the hyphal growth on solid media, particularly when GlcNAc was used as the sole source of carbon. The hyphal-growth inhibition was also significant in the *qcr7Δ/Δ* mutant under liquid-induced medium with GlcNAc as the sole carbon source, but not with other carbon sources. In addition, the activity of hyphae-related hydrolases decreased.

A global transcriptional approach was adopted to investigate which genes influence these phenotypic changes. Within the GO category “biological processes”, selected pathways associated with crucial virulence factors, such as those gene involved in filamentous growth, biofilm formation, hyphal growth, and cell adhesion, were downregulated, as well as genes assigned to transport, carbohydrate metabolic processes, and lipid-metabolic processes. Those with significant differential changes were found to be involved in carbohydrate transport on the basis of the functional analysis of the downregulated genes. Although partially related genes were upregulated to compensate for the effect, the effect was insignificant, which indicated that QCR7 deletion weakened the carbohydrate-use ability of the *C. albicans*. For the *C. albicans*, it was necessary to coordinate the use of sugars as energy sources with the production of active sugars (uridine diphosphate (UDP)-glucose, guanosine diphosphate-mannose, and UDP-N-acetylglucosamine), which combine to synthesize major cell-wall polysaccharides (glucan, mannose, and chitin) for the biosynthesis of new cell walls (Piškur and Compagno, 2014). However, as expected, the QCR7 deletion affected the integrity of the cell wall, which was consistent with the transcriptional response of the *qcr7Δ/Δ* mutant. Alternatively, for the GO cellular components, the GO terms associated with the membrane and cell wall were significantly enriched in the downregulated gene sets. Notably, the mutant exhibited hypersensitivity to various cell-wall stressors, such as CFW, caspofungin, and CR.

According to the comparative transcriptome analysis and several pathogenic phenotypes of the *qcr7Δ/Δ* mutant, these data revealed that the QCR7 deletion had the most significant effect on biofilm formation. In *C. albicans*, a core complex regulatory network is critical for biofilm formation. Here, we showed that the six core biofilm-regulatory-factor mutants exhibited significantly downregulated expression of QCR7.We then increased the expression level of QCR7 in these mutants and performed a comparison of the biofilm-forming ability, showing that increased expression levels of QCR7 partially restored the biofilm defects caused by the NDT80 knockdown. Our findings provide additional support for the role of QCR7 in biofilm formation regulated by NDT80. In addition, to better understand whether the significant downregulation of biofilm-formation-related genes in the *qcr7Δ/Δ* mutant was the key factor affecting the virulence factors, we overexpressed five significantly downregulated biofilm-formation-related genes in the *qcr7Δ/Δ* mutant background, showing that the biofilm phenotype of the *qcr7Δ/Δ* mutant was partially reversed after the overexpression of four genes, while the overexpression of one gene further attenuated the biofilm phenotype of the *qcr7Δ/Δ* mutant. Intriguingly, YWP1 is a yeast-forming cell-wall protein, which is known to inhibit biofilm formation (Granger et al., 2005). HYR1 is a GPI-anchored cell-wall protein, which is expressed during hyphal development. There are studies that prove that this mutant may have a moderately attenuated biofilm phenotype (Dwivedi et al., 2011). However, the biofilm-mass formation of *qcr7Δ/Δ* mutant was increased by the overexpression of *YWP1*,while the biofilm-deficient phenotype of the *qcr7Δ/Δ* mutant was more pronounced through the overexpression of *HYR1*. Previous studies identified *HWP1, XOG1*, and *SAP6* as biofilm-massociation genes for managing *C. albicans* virulence (Nobile et al., 2006; Taff et al., 2012; Winter et al., 2016). The increased expression of these three genes in the *qcr7Δ/Δ* mutant background could partially rescue the biofilm phenotype. The hyphal growth on solids of these five overexpressing strains agrees with observations on biofilm formation, according to which *HWP1, YWP1, XOG1*, and *SAP6* overexpression partly but significantly reversed the *qcr7Δ/Δ* mutant phenotype, and yet *HYR1* overexpression additively attenuated pathogenic phenotypes. A similar phenomenon was also observed in the experiment on carbon-source utilization. Taking into account all these factors, we may reasonably conclude that the deletion of QCR7 had the most significant effect on the cell surface, thus affecting a variety of virulence factors. The increase in the expression level of a series of gene-encoding cell-surface proteins resulted in the restoration of the corresponding virulence phenotype and carbon-source utilization. In summary, we propose a model (Figure 7) for the role of QCR7 in promoting biofilm formation in *C. albicans*. In our study, we found that QCR7 plays a key role in a range of biological functions and structural components of *C. albicans*, and that these have an impact on the formation of *C. albicans* biofilms. The deletion of QCR7 had significant effects on carbon utilization, cell-surface gene expression, biofilm formation, and hyphal-growth maintenance, and the molecular-mechanism exploration revealed a close correlation between these phenotypic changes. In an in-depth study of the effect of QCR7 on biofilm formation, it was found that QCR7 was regulated by NDT80 and regulated cell-surface-related genes, mitochondrial function, etc. which reduced the formation of dense biofilm matrix.

**Figure 7.**
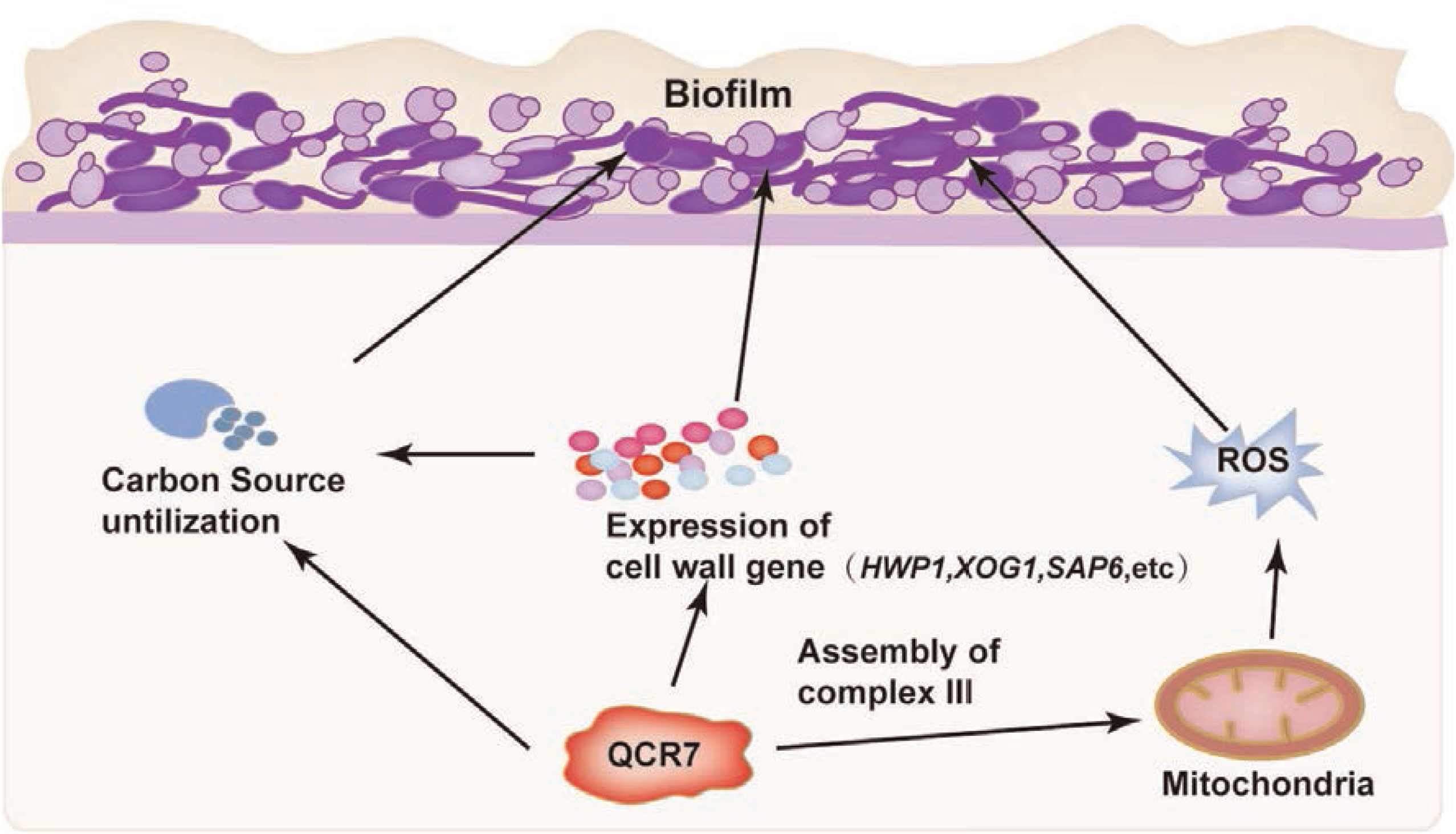
A model of the role of QCR7 in regulating multiple biological processes affecting biofilm formation. The important role of QCR7 in carbon utilization, affecting cell-surface gene expression and mitochondrial function, provides important clues as to the mechanisms through which QCR7 affects biofilm formation. Defective biofilm formation mainly attributed to the downregulation of multiple-cell-surface genes was confirmed experimentally, while the expression of these genes was also identified as critical in influencing carbon source utilization. The cross-linking of multiple pathways provides a possible mechanism through which QCR7 regulates dense-biofilm-matrix production through the carbon-source environment of the host and the regulation of cell-surface gene expression.

## 5. Conclusion

Our study suggests that QCR7 affects carbon absorption and use, thus influencing cell-surface function, reducing invasion, and, finally, infecting host tissues.

## 6. Data availability statement

Raw sequencing data were submitted to the National Center for Biotechnology Information database with the bioproject ID PRJNA802957.

## 7. Ethics statement

The ethics committee of the Nanchang University approved this study (approval number: NCUSYDWFL-201035).

## 8. Author contributions

LBZ, YCH, JJT, and XTH designed the study. LBZ, YCH, JJT, NYH, YPZ, JZC, QL, and YLC performed the experiments and data analyses and wrote the original draft. LBZ and XTH edited the manuscript. All the authors have read and approved the final version of the manuscript.

## 11. Acknowledgments

We thank the Nanchang University Medical School for providing the space and equipment necessary to conduct this study.

## Supplementary Information

**Figure S1.**
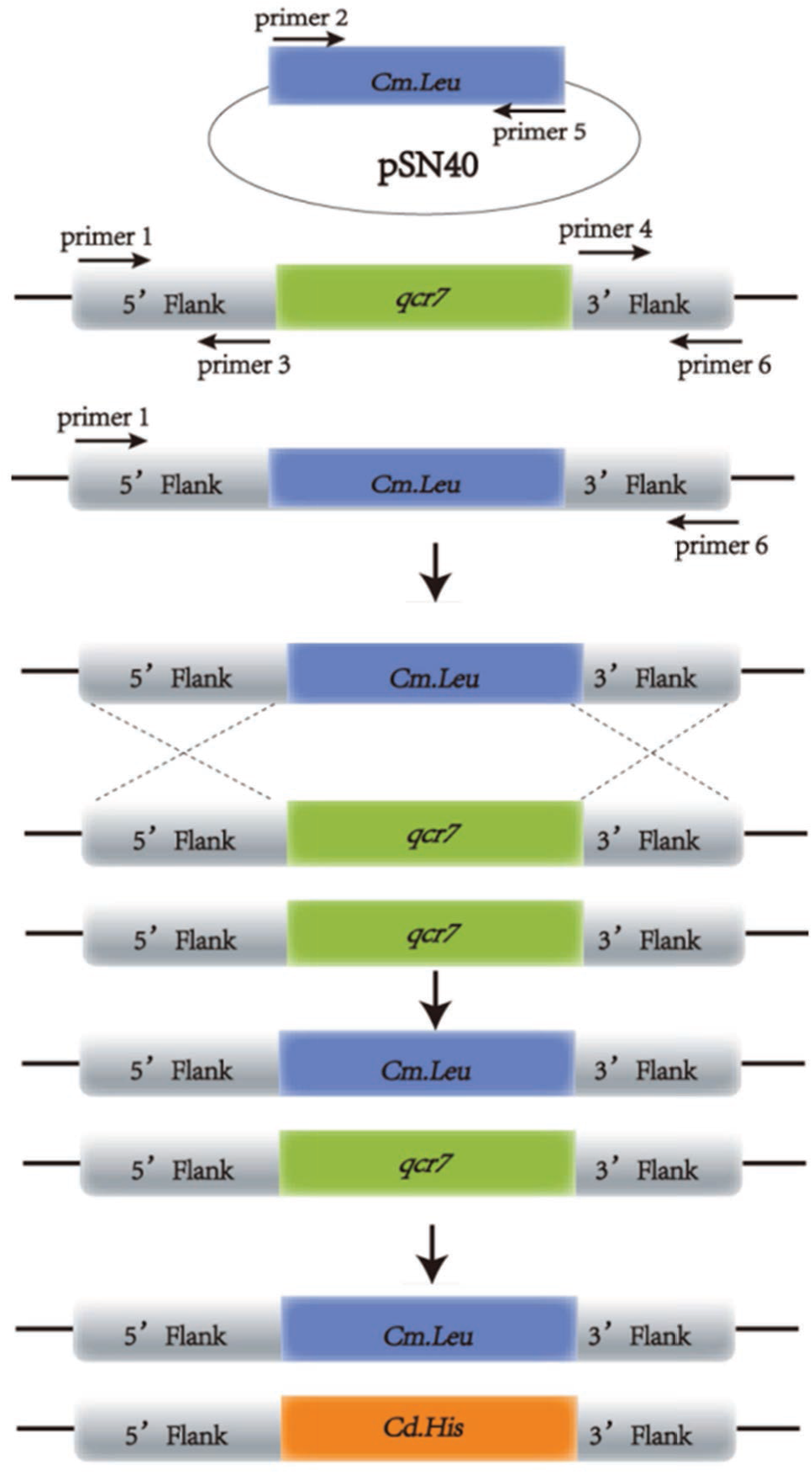
Schematic diagram of the gene-knockout strategy using PCR-based homologous recombination. Primers used in the work were: Amplified *LEU2/HIS1* cassette (Universal primer 2+Universal primer 5), amplified *QCR7* gene 5′fragments (primer 1+primer 3), amplified *QCR7* gene 3′fragments (primer 4+primer 6).

**Figure S2.**
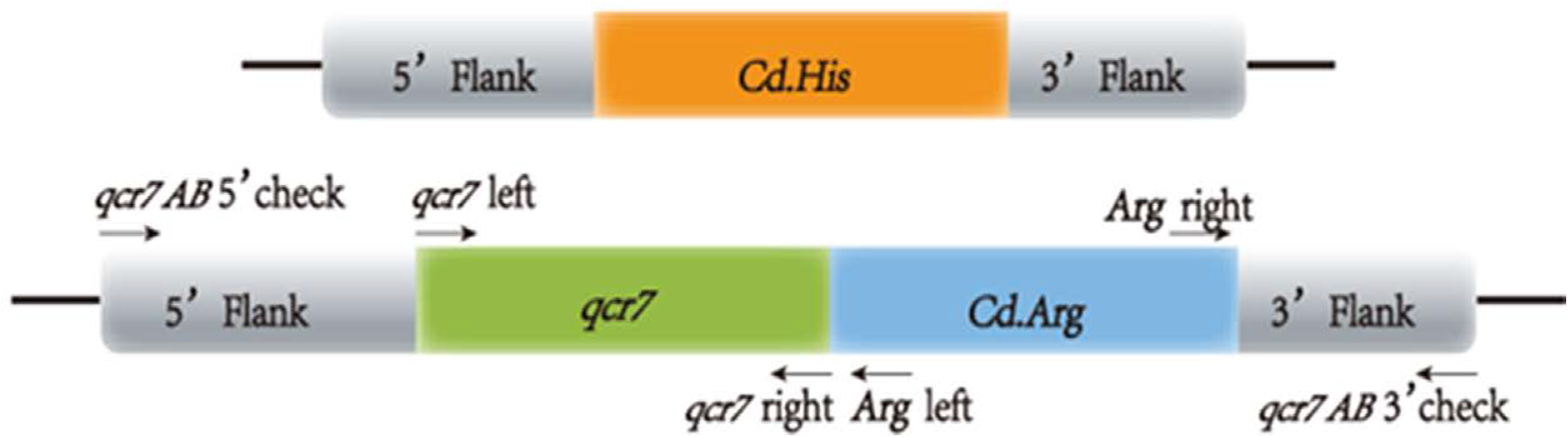
Schematic diagram of the gene-reconstitution strategy using complementation via PCR-based homologous recombination. Primers used in the work were: Amplified the entire *QCR7* ORF with flanking upstream (primer 1+primer 13), amplified *ARG4* cassette (Universal primer 2+Universal primer 5).

**Figure S3.**
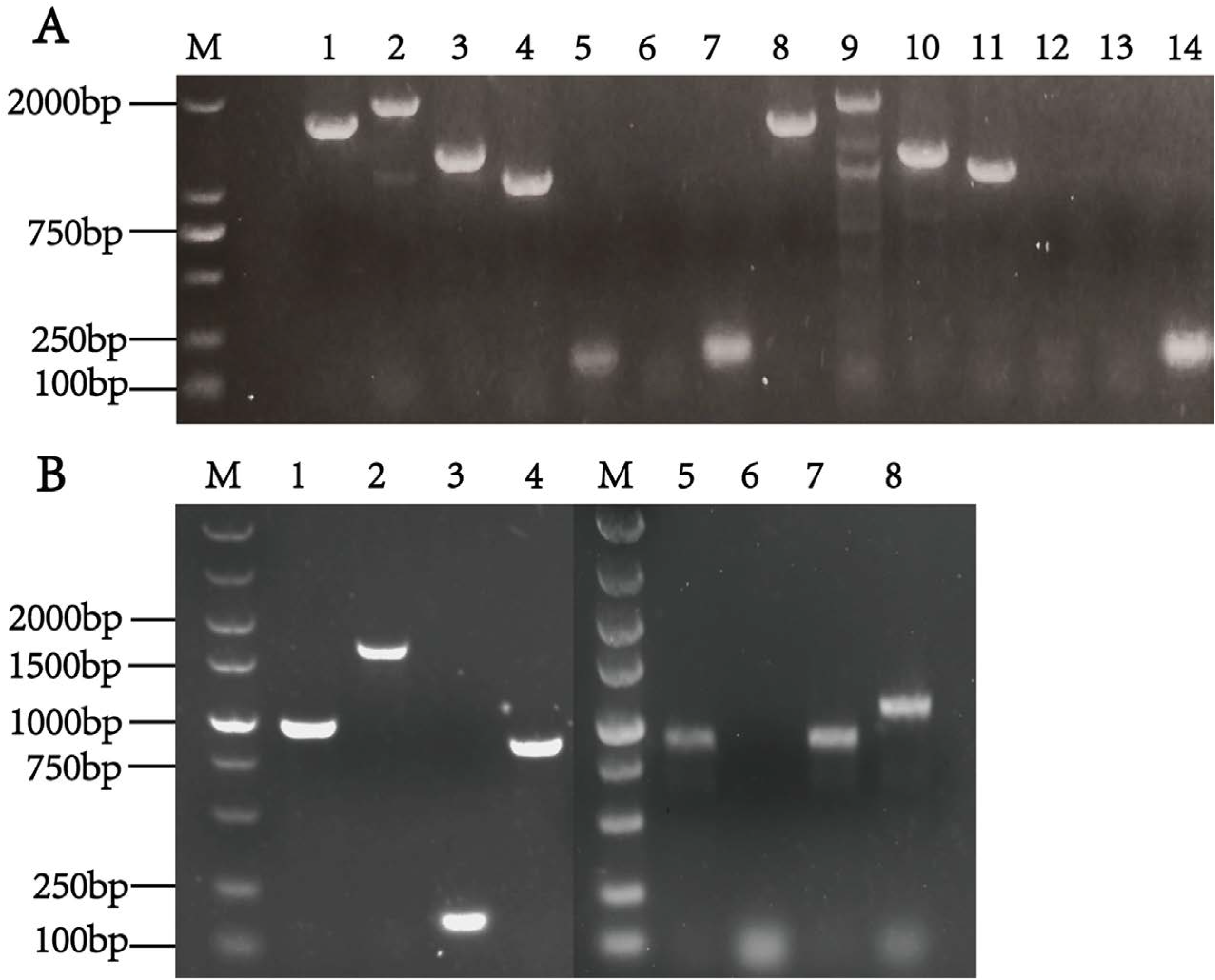
Identification of the *QCR7* knockout and reconstituted strains. (A) PCR-based identification of the *QCR7* knockout strains of *C. albicans*, *QCR7* upstream check, *LEU2* left and right, *QCR7* downstream check, and *QCR7* upstream check, *HIS1* left and *HIS1* right, and *QCR7* downstream check, were used to verify whether the marker gene was transformed into *C. albicans*. The lane8 are stripe at 1,575 bp for *qcr7Δ/Δ* with primer *Qcr7* upstream check and *LEU2* left, lane9 are stripe at 1,853 bp for *qcr7Δ/Δ* with primer *Qcr7* downstream check and *LEU2* right, lane10 are stripe at 1,293 bp for *qcr7Δ/Δ* with primer *Qcr7* upstream check and *HIS1* left, lane11 are stripe at 1,241 bp for *qcr7Δ/Δ* with primer *Qcr7* downstream check and *HIS1* right, lane12 are without stripe for *qcr7Δ/Δ* with primers *QCR7* left and right, lane13 are without stripe for H2O as a template with primers *QCR7* left and *QCR7* right, and lane14 are stripe at 169 bp for WT with primers *QCR7* left and *QCR7* right. The band sizes were in line with expectations. (B) PCR-based identification of the gene-reconstituted strains. Using the QCR7AB genome as a template, the primers QCR7AB 5′ check (*QCR7* upstream check) and *QCR7* right, QCR7AB 5′ check (*QCR7* upstream check) and *ARG* left*, QCR7* left and *QCR7* right, *QCR7* left and *ARG* left, and *ARG* right and *QCR7* AB 3′ check (*QCR7* downstream check). The question of whether *QCR7* complement transformation component was integrated in a site-specific manner was verified. Lane1 are stripe at 987 bp for QCR7AB with primer *QCR7* upstream check and *QCR7* right, lane2 are stripe at 1,634 bp for QCR7AB with primer *Qcr7* upstream check and *ARG* left, lane3 are stripe at 169 bp for QCR7AB with primers *QCR7* left and right, lane4 are stripe at 816 bp for QCR7AB with primer *Qcr7* left and *ARG* left, lane8 are stripe at 1093 bp for QCR7AB with primer *ARG* right and *qcr7* AB 3′ check (*QCR7* downstream check). The band sizes met the expectations. These results suggest that the *QCR7* nonmutant and reconstituted strains were constructed successfully.

**Figure S4.**
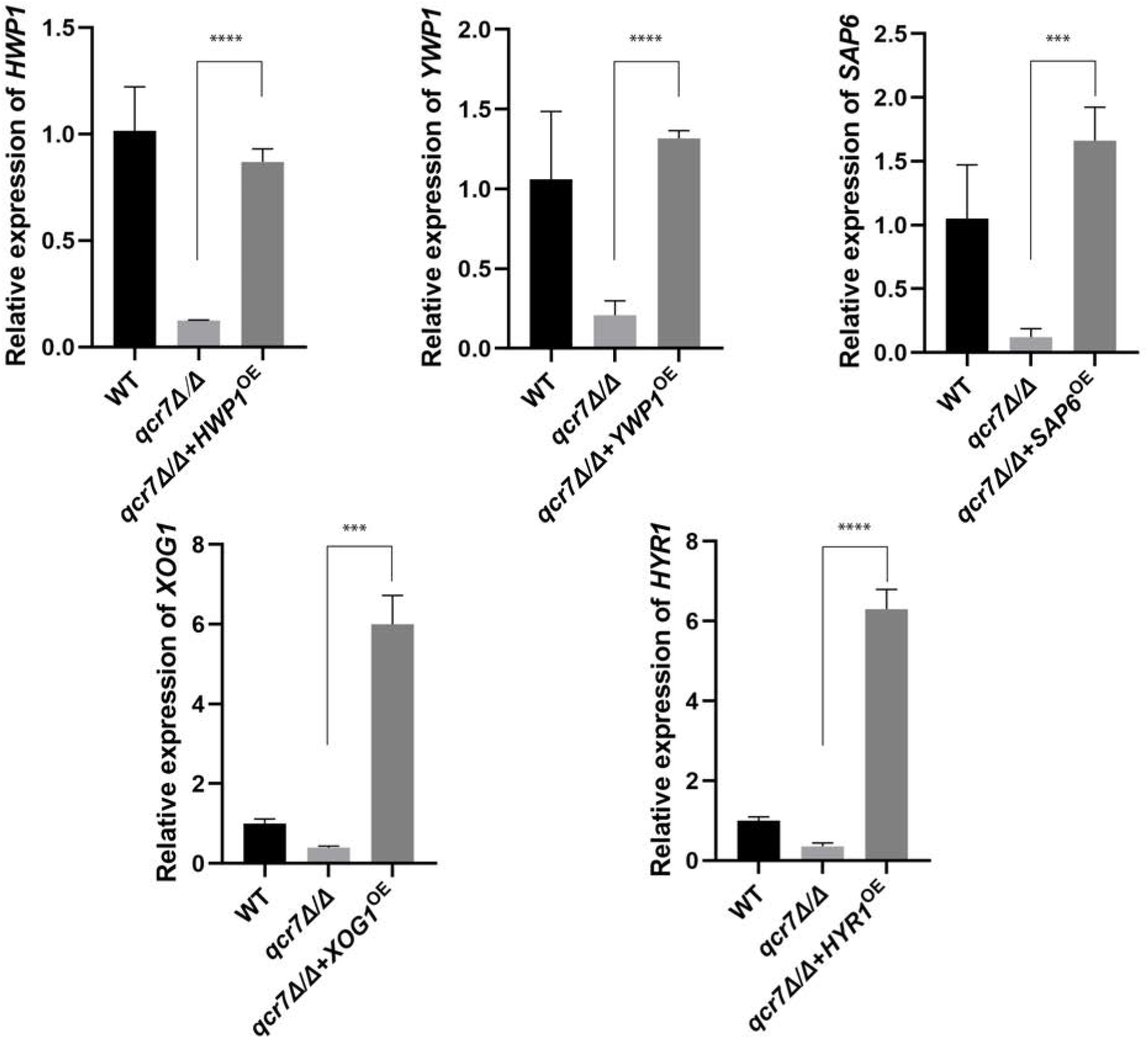
The construction and integration of inserts of a wild-type copy of the cognate gene into NEUT5L locus results in the overexpression of related genes in genetic background. RT-qPCR analysis of relative transcript levels of listed genes in the wild-type, mutant, and overexpression strains.

**Figure S5.**
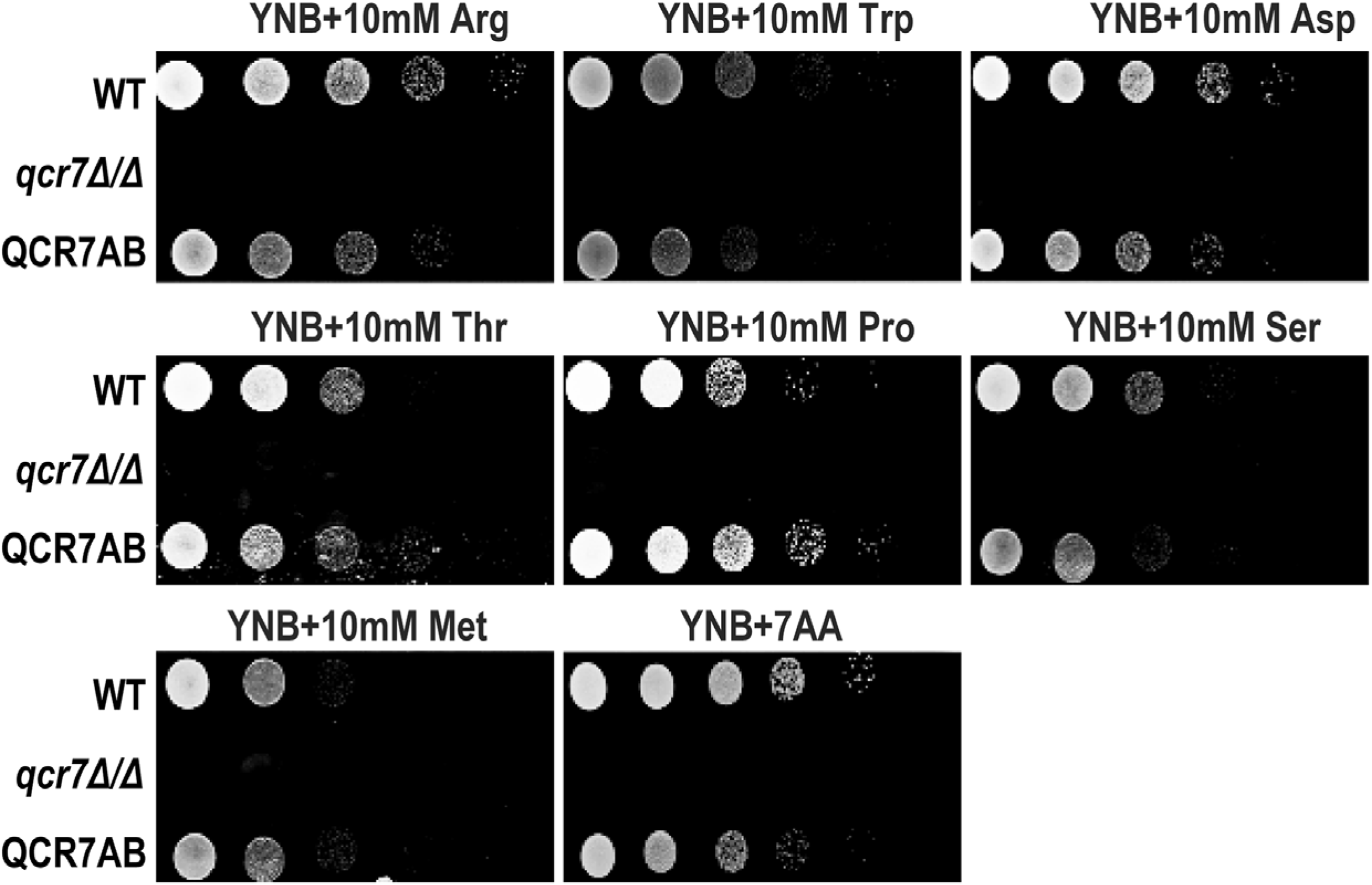
*QCR7* deletion does not lead to the use of amino acids as carbon sources. Strains were cultured overnight in YPD, washed, and serially diluted with PBS. Different concentrations (5μl of 10^7^ cells/ml to 10^3^ cells/ml) of WT, *qcr7Δ/Δ*, and QCR7AB were spotted on solid YNB media containing 5g/L (NH4)_2_SO_4_ and supplemented with several amino acids as the sole carbon sources. Plates were photographed after incubation at 30°C for 2 days.

**Figure S6.**
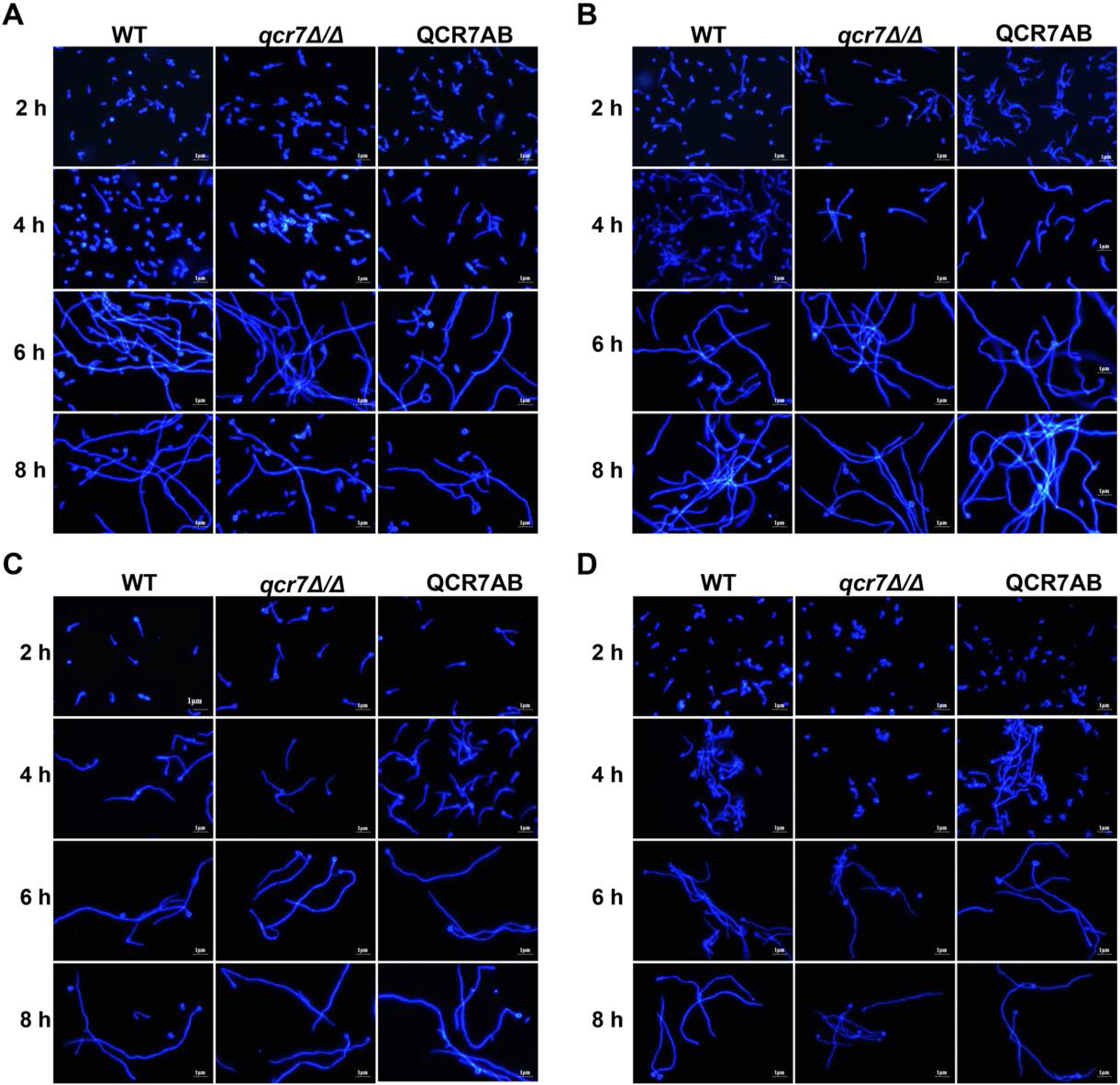
*QCR7* deletion is not evident in defective hyphae cultured using fermentable carbon source. The wild type, *qcr7Δ/Δ*, and QCR7AB were cultured overnight in liquid YPD at 30°C, followed by washing and PBS dilution to OD_600 nm_ = 0.1, after which the mixture was resuspended in Spider medium supplemented with glucose(A), mannitol(B), sucrose (C) or maltose (D); the incubation was continued at 37°C. The hyphal morphologies were observed under a fluorescence microscope (Olympus, Japan). Scale bars are 1 μm.

**Figure S7.**
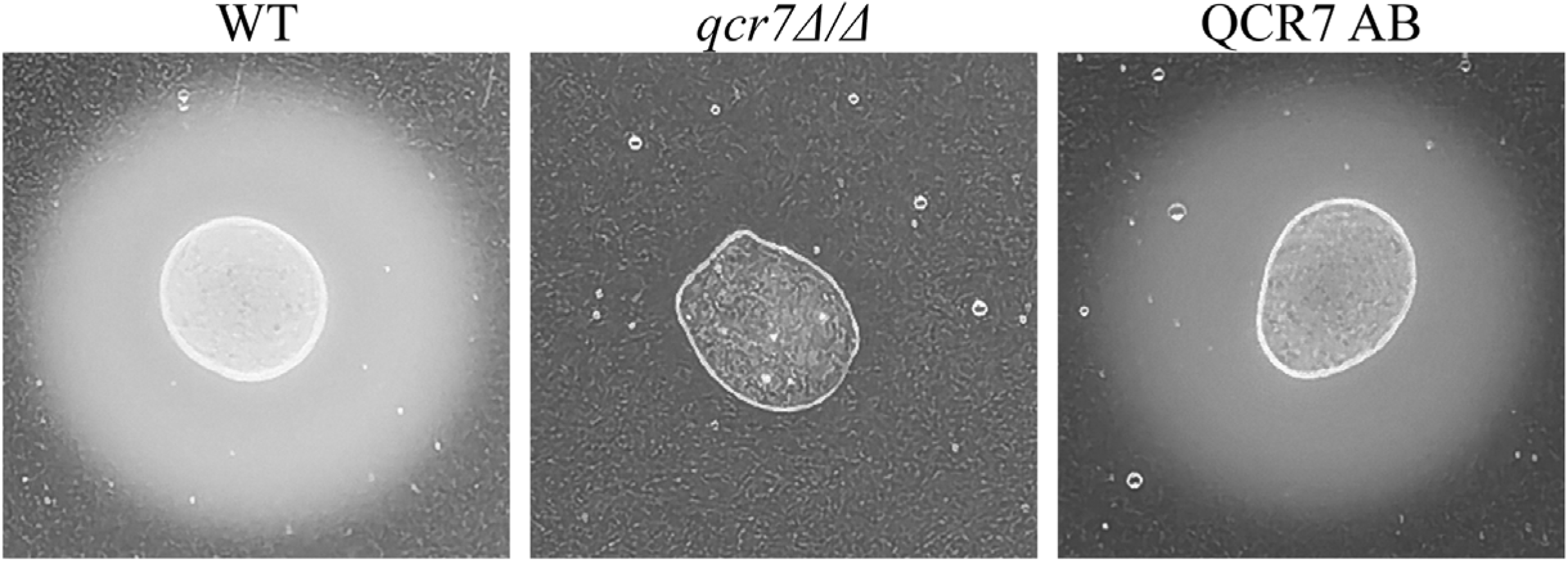
Wild-type and *qcr7Δ/Δ* strains exhibit different secreted aspartyl protease activities. Sap-activity assays were performed using a medium with BSA as the sole nitrogen source. Then, 1 × 10^5^ cells suspended in water were spotted on YNB-BSA agar plates and incubated at 37°C for 5 days. The size of white halo rings indicates the activity levels.

**Figure S8.**
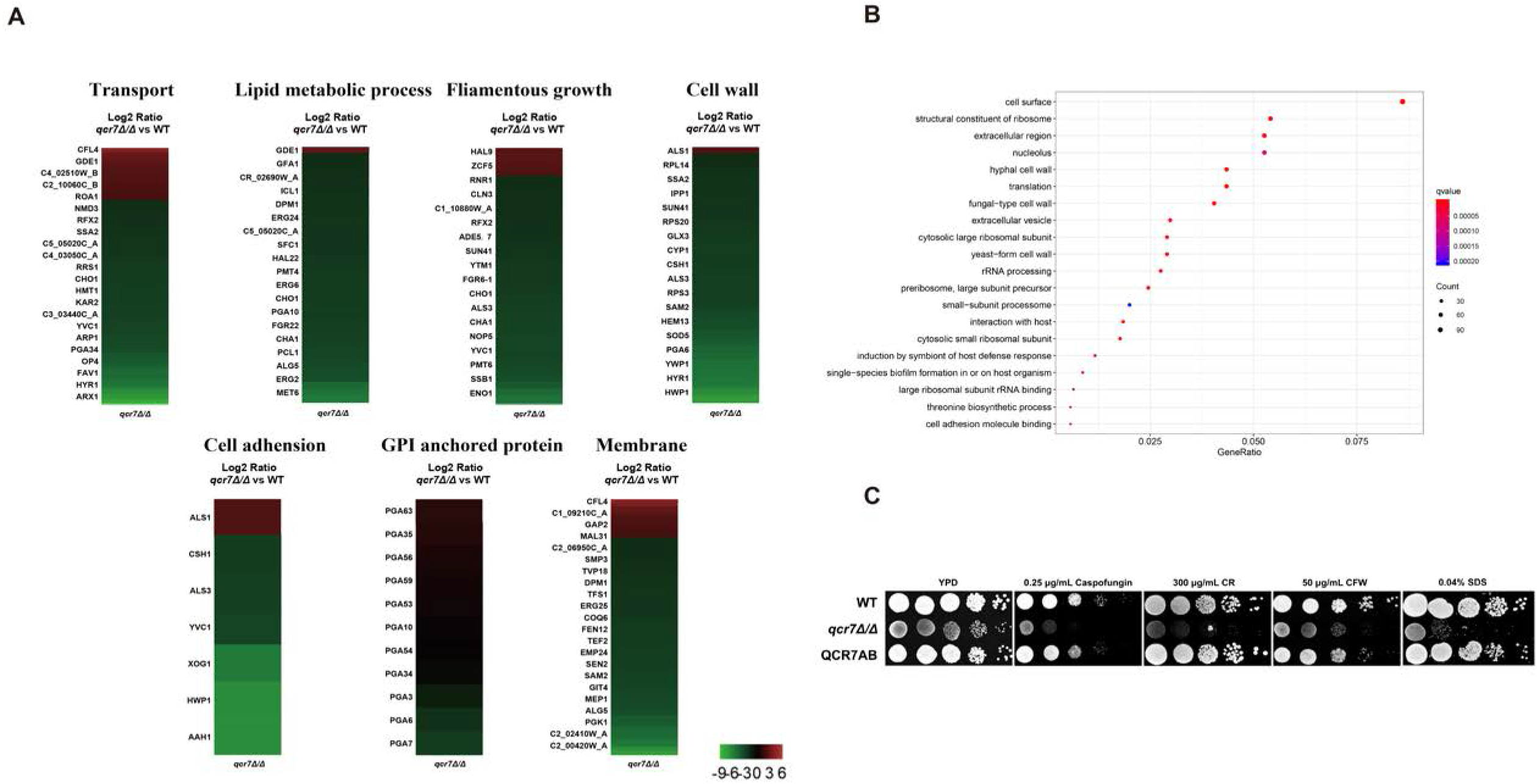
QCR7 is required for *C. albicans* cell-wall maintenance and expression of cell-surface genes. (A) A heatmap displaying the upregulated and downregulated genes that associate partly with the GO category “cellular component” or “biological processes.” Fold enrichment of these genes is shown in Table S3. (B) KEGG-pathway-enrichment analysis (p < 0.05). (C) Strains were cultured overnight YPD, after which they were washed and serially diluted with PBS. Different concentrations (5μl of 10^7^ cells/ml to 10^3^ cells/ml) of strains were then spotted on YEP containing cell-wall stressors (300 μg/ml CR, 50 μg/ml CFW, and 0.25 μg/ml caspofungin, 0.04%SDS).

**Figure S9.**
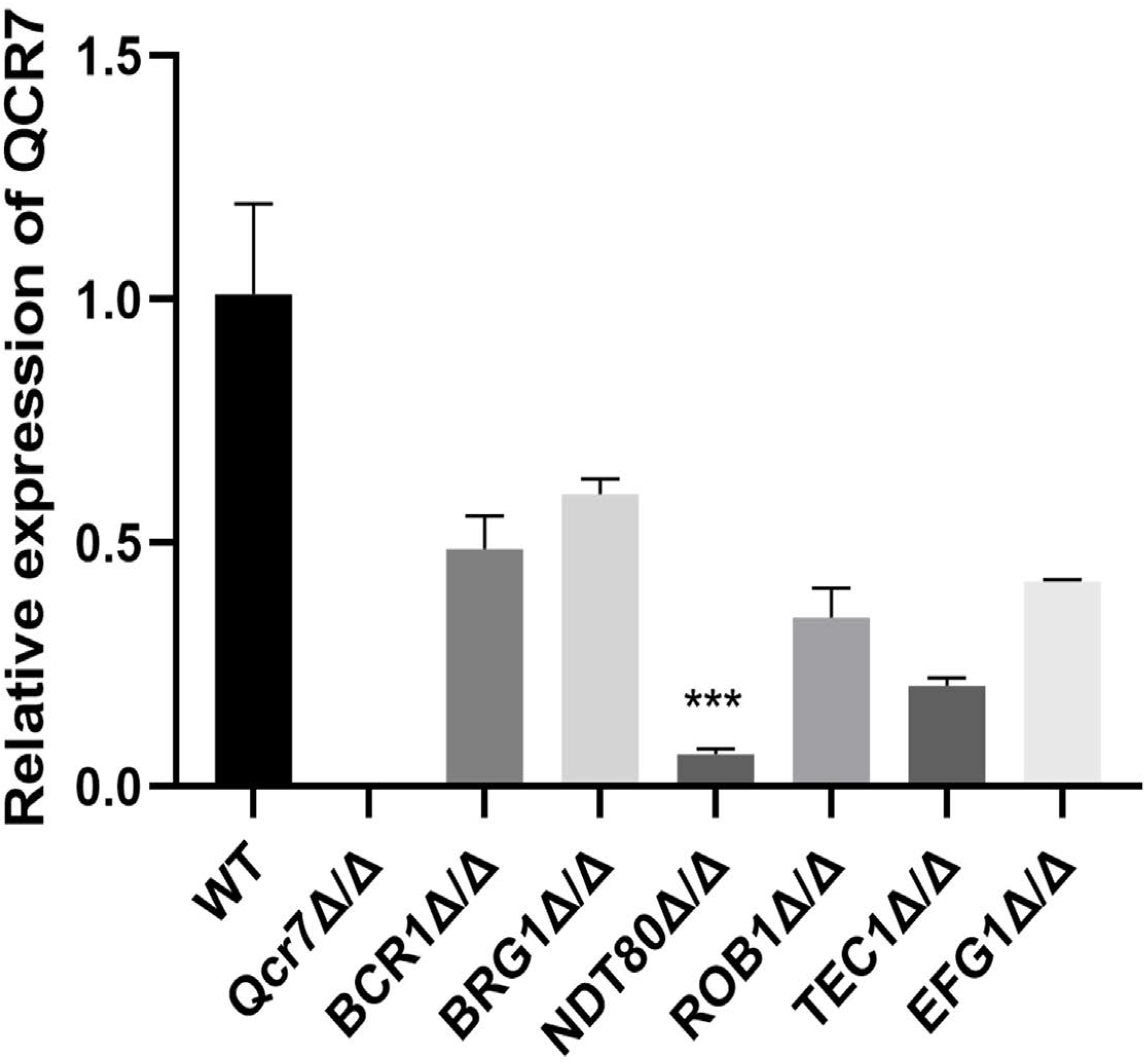
QCR7 plays a key role in biofilm formation. Deletion of *BRG1, ROB1, NDT80, TEC1*, or *EFG1* all had an effect on *QCR7* expression.

**Figure S10.**
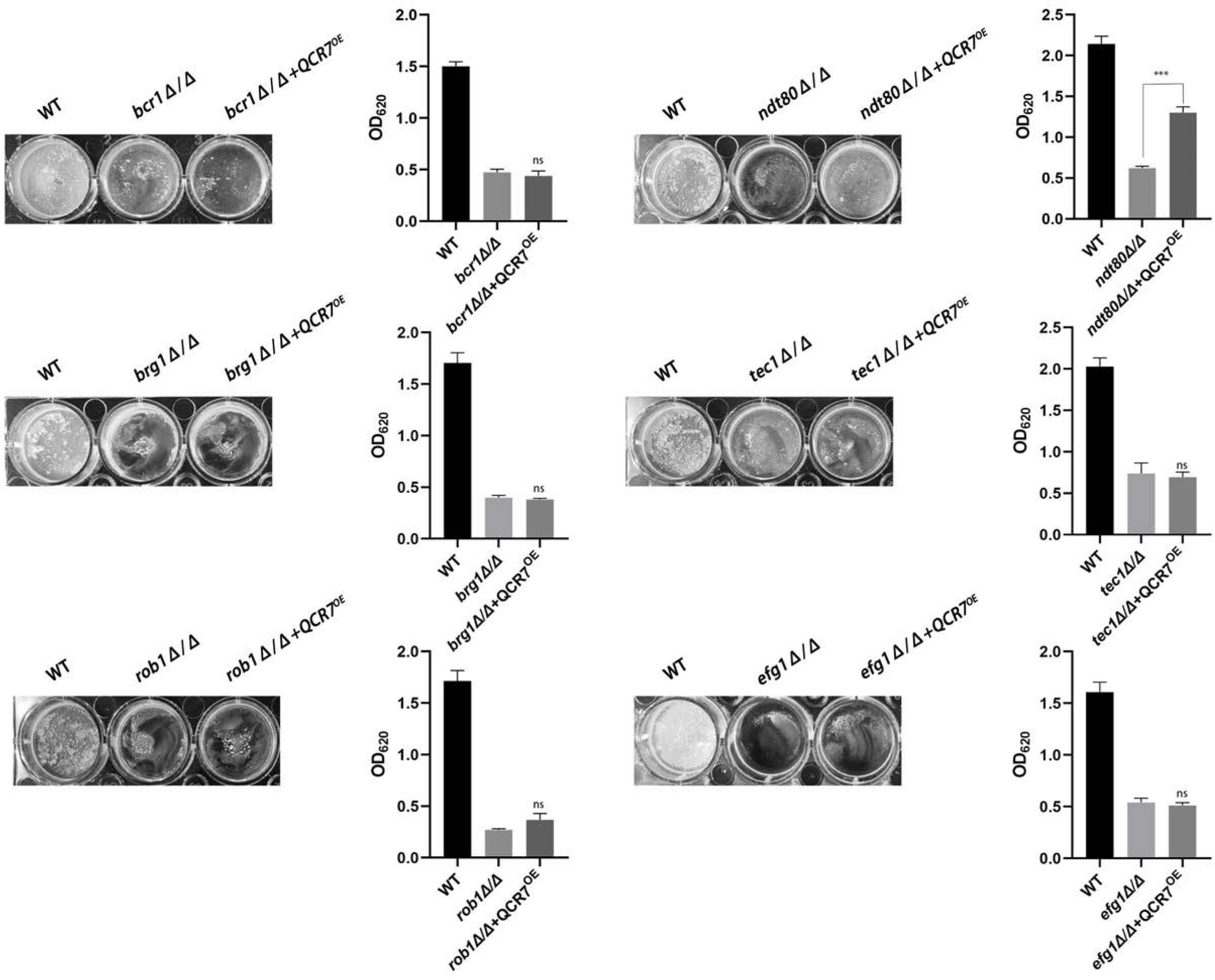
NDT80 regulation of biofilm formation is partially dependent on QCR7. Biofilm formation (48h, 37°C) of strains was profiled and statistically compared to reference strain *C. albicans* SN250.

## Tables

**Table S1.**
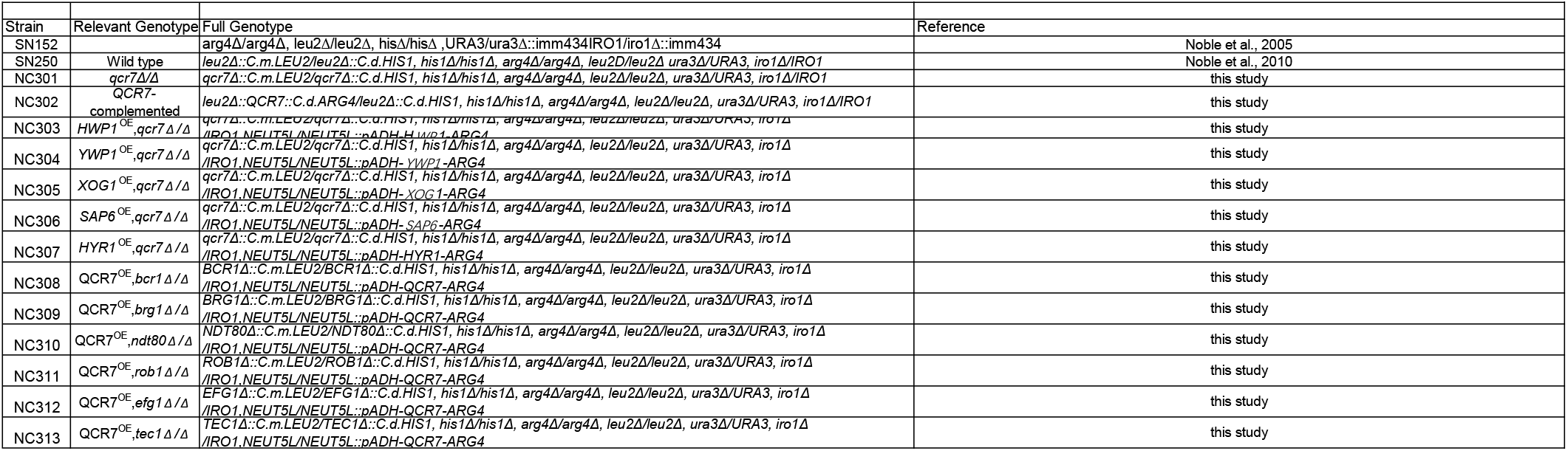
Strains used in this study.

**Table S2.**
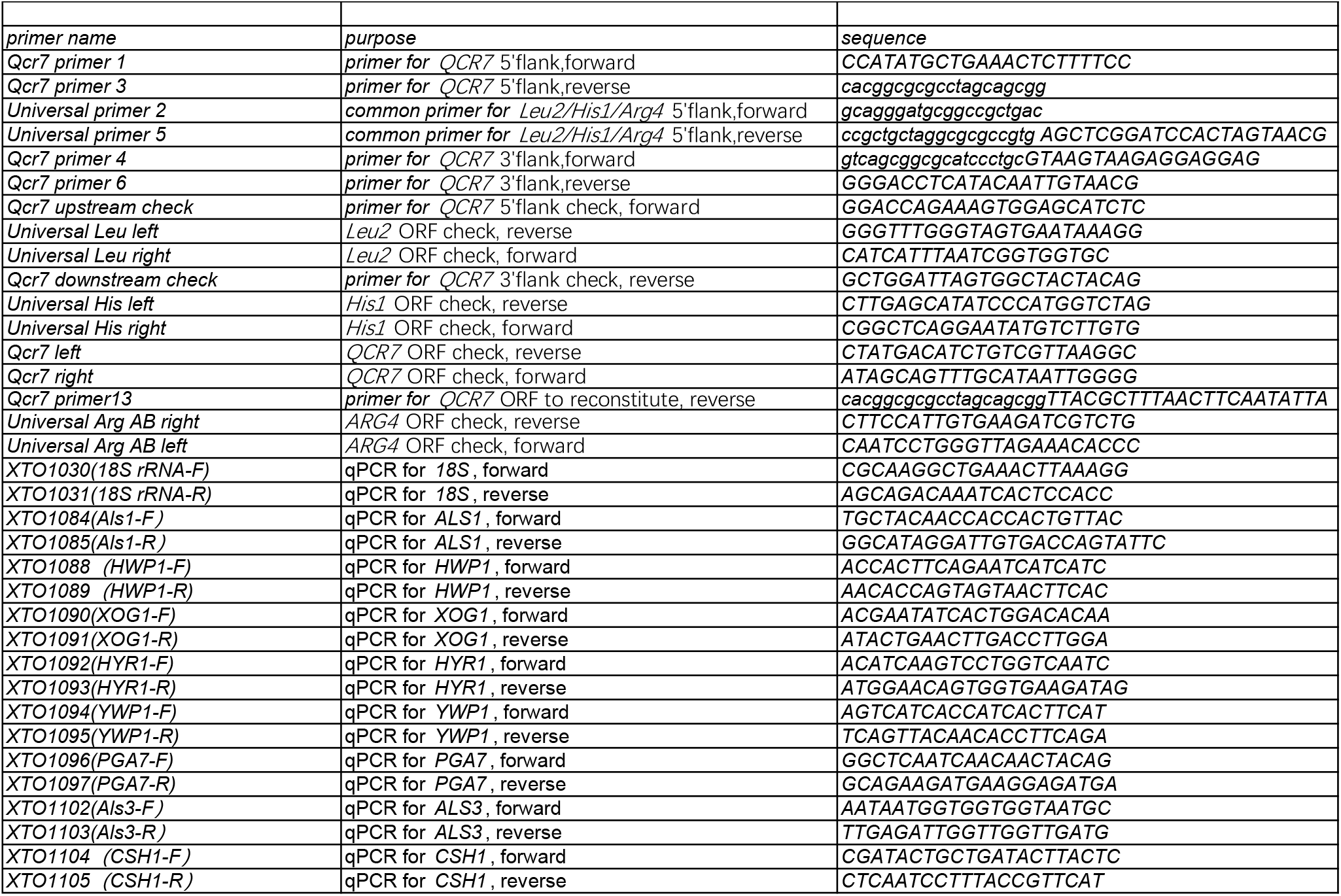
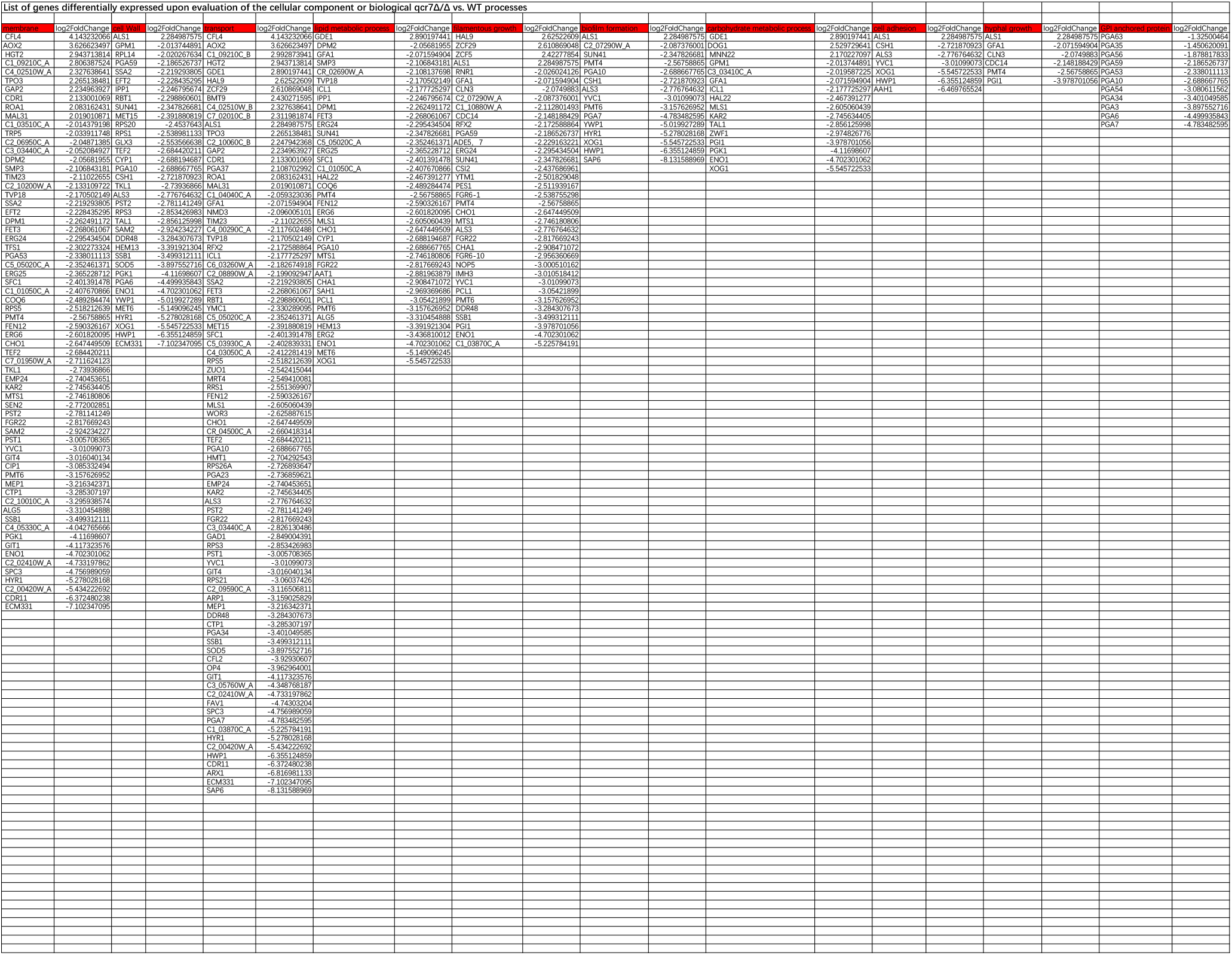
Primers used in this study.

Table S3 List of genes differentially expressed upon the evaluation of cellular components or biological *qcr7Δ/Δ vs*. WT processes

